# Space-for-time substitution reveals partial transferability of marine biodiversity-temperature relationships

**DOI:** 10.64898/2026.08.19.745747

**Authors:** Marina Costa Rillo, Lea Möller, Lukas Jonkers, Julian Merder, Helmut Hillebrand

**Affiliations:** GeoZentrum Nordbayern, FAU - Friedrich-Alexander University Erlangen-Nürnberg, Erlangen, Germany; ICBM - Institute for Chemistry and Biology of the Marine Environment, Carl Von Ossietzky Universität Oldenburg, Oldenburg, Germany; HIFMB - Helmholtz-Institute for Functional Marine Biodiversity, University of Oldenburg, Oldenburg, Germany; MARUM - Center for Marine Environmental Sciences, University of Bremen, Bremen, Germany; Department of Environmental Science, Aarhus University, Roskilde, Denmark; AWI - Alfred Wegener Institute, Helmholtz-Centre for Polar and Marine Research, Bremerhaven, Germany

**Keywords:** biodiversity change, climate change, conservation paleobiology, hindcasting, microfossils, paleoecology, prediction

## Abstract

Forecasts of biodiversity responses to climate change often rely on space-for-time substitution, in which spatial biodiversity-climate relationships are used to predict biodiversity change through time. Yet this approach is rarely tested directly because long-term biodiversity time series are scarce. Here, we combine global modern and fossil assemblage data of planktonic foraminifera with site-specific sea-surface temperature reconstructions to compare biodiversity-temperature relationships across space and time. Spatial and temporal compositional turnover models showed similar slopes but consistently different intercepts, with spatial models predicting higher turnover across the full temperature gradient. Restricting the spatial comparison to the environmental domain of individual fossil time series reduced, but did not eliminate, this intercept mismatch. For alpha diversity, spatial models more closely recovered the temporal biodiversity-temperature relationship than for compositional turnover. Thus, for the timescales studied here, space-for-time substitution captures the direction of biodiversity change but not its magnitude through time.

## 1 | INTRODUCTION

Ecologists face the challenge of predicting how biodiversity will change in response to anthropogenic climate change. Such forecasts are widely used in conservation biology (Dawson et al., 2011; Mouquet et al., 2015), yet most ecological time series lack sufficient climatic variation to validate them (Dornelas et al., 2018; Adler et al., 2020) or can only estimate temperature effects on the initial rate of biodiversity change (Pinsky et al., 2025). A widely used approach is space-for-time substitution (Pickett, 1989; Lovell et al., 2023), which assumes that spatial relationships between climate and biodiversity can be used to predict biodiversity dynamics through time under changing climates. Its appeal lies in the idea that spatial gradients can capture a broader range of environmental variation than is often observed in time-series data, allowing ecological responses estimated across space to be extrapolated to temporal change. Whether spatial biodiversity-climate relationships can accurately predict temporal dynamics remains rarely tested (Lovell et al., 2023; Kharouba and Williams, 2024), and the fossil record provides one of the few ways to directly evaluate this assumption by comparing spatially derived relationships with past temporal change (i.e., hindcasting).

Space-for-time substitution underpins widely used biodiversity forecasting approaches, including species distribution models (Elith and Leathwick, 2009; Guisan and Thuiller, 2005; Hodapp et al., 2023) and models of phenological responses (Ford et al., 2016; Zografou et al., 2020), genetic patterns (Wogan and Wang, 2018), and ecosystem functions (Frauendorf et al., 2020). Despite this broad use, empirical tests of space-for-time substitution have yielded mixed results. Using Quaternary pollen assemblages from North America, Blois et al. (2013) showed that spatial models captured temporal turnover, particularly when spatial and temporal climate variation were comparable. This work has since been widely cited as support for the use of space-for-time substitution in biodiversity forecasting. Other tests have similarly found good transferability (e.g. Illán et al. 2014; Guerin et al. 2012), but many have not: spatial relationships have failed to predict temporal change in birds (La Sorte et al., 2009), glacial-refugia distributions Veloz et al. (2012); Davis et al. (2014); Worth et al. (2014), and tree growth, where spatial and temporal responses to temperature pointed in opposite directions (Perret et al., 2024; Klesse et al., 2020). This inconsistency appears to depend on the taxa, ecological level, and timescale being modelled (Wogan and Wang, 2018; Kharouba and Williams, 2024; Lovell et al., 2023; Evans et al., 2025).

Space-for-time substitution has not been directly tested in marine systems, despite the availability of rich sedimentary archives. Marine microfossils have long been used in paleoenvironmental reconstructions (Kucera, 2007), which rely on the same space-time substitutability assumption but solve the reverse problem: inferring past environments from assemblages rather than predicting assemblages from environments. Planktonic foraminifera are globally distributed unicellular eukaryotes and provide a particularly suitable system for such a space-time validation. Their assemblage composition and diversity gradients are strongly structured by temperature (Rutherford et al., 1999; Fenton et al., 2016; Morey et al., 2005; Rillo et al., 2022); species’ thermal niches appear to have remained broadly stable over the last 800,000 years (Antell et al. 2021, but see Ying et al. 2024); and abundant assemblage data are available from both modern surface-sediment and fossil sediment-core records across broad spatial and temporal scales (Siccha and Kucera, 2017; Strack et al., 2022; Fenton et al., 2021). In addition, fossil assemblages integrate community change over hundreds to thousands of years (Jonkers et al., 2019), reducing the likelihood that short-term transient dynamics dominate the temporal signal (Adler et al., 2020).

Here, we use globally distributed ocean-floor surface sediment (modern assemblage) and sediment core (fossil time series) data of planktonic foraminifera spanning the past 250 thousand years, together with corresponding site-specific sea-surface temperature estimates from Mg/Ca and alkenone proxies (Fig. 1). We test for space-time equivalence (Lovell et al., 2023) by comparing the slope and intercept of biodiversity-temperature relationships estimated in space and through time. We focus on two dimensions of biodiversity: beta diversity, measured as compositional turnover, and alpha diversity, measured as effective numbers of species. Because planktonic foraminiferal assemblages show strong temperature dependence both spatially and temporally, we expect spatial models to recover temporal biodiversity change (Wogan and Wang, 2018). Our results, however, show that this recovery is partial: spatial and temporal models yield similar slopes – indicating a shared direction and rate of change in the biodiversity-temperature relationship – but spatial models consistently predict a higher intercept, overestimating the magnitude of turnover across the full temperature gradient.

**F I G U R E 1.**
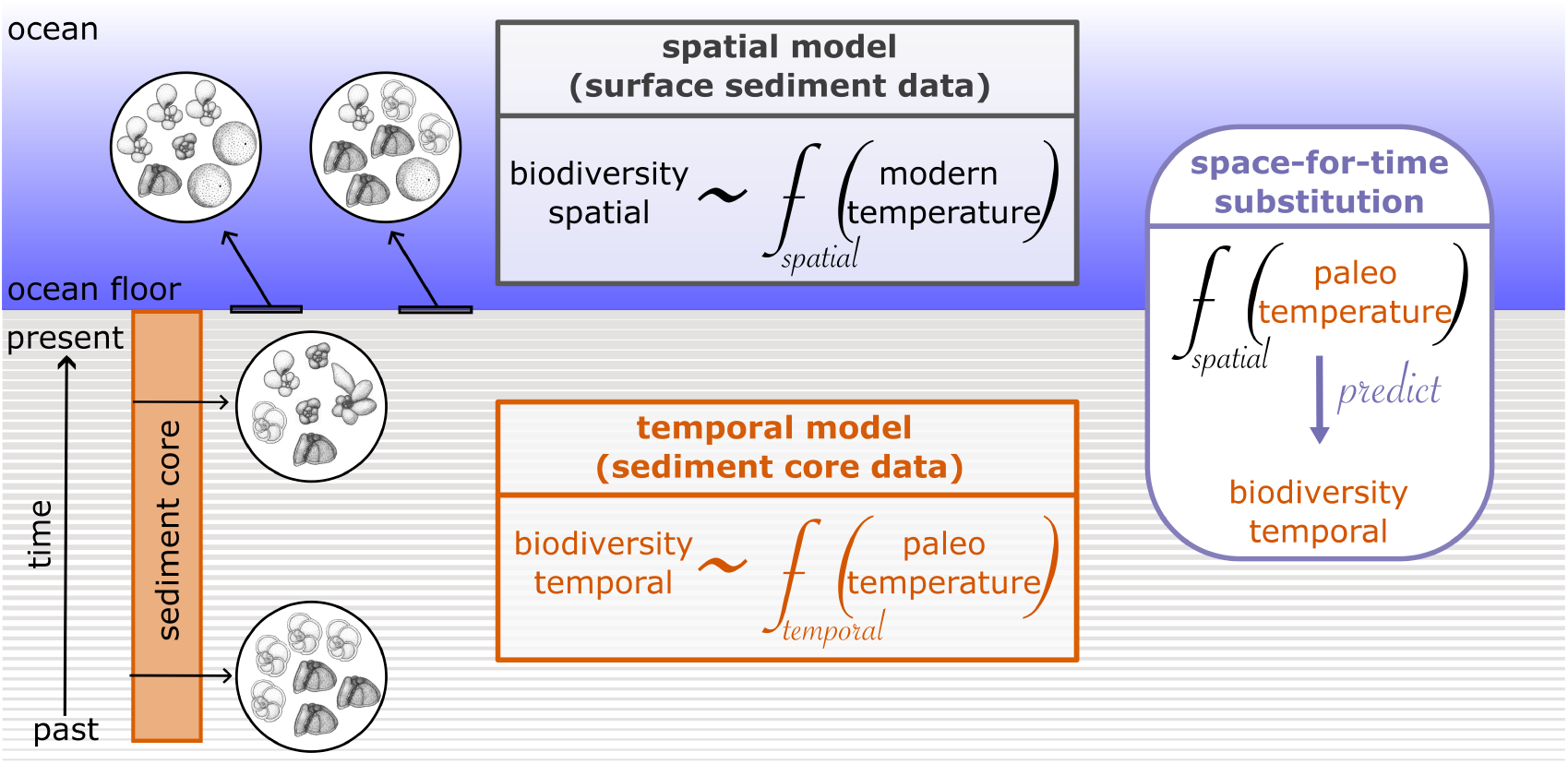
Conceptual framework for testing space-for-time substitution in marine plankton biodiversity. Modern surface sediment assemblages and associated present-day sea-surface temperatures are used to estimate spatial biodiversity-temperature relationships across the ocean. Fossil assemblages and corresponding site-specific paleo-temperature reconstructions from sediment cores are used to estimate temporal biodiversity-temperature relationships through time. Comparing the two provides a direct hindcast test of whether biodiversity-temperature relationships inferred from spatial variation predict biodiversity change through time. Specimens’ drawings from Parker (1962). The “f” corresponds to the biodiversity-temperature relationship (function) and was fitted with spatial (*f*_*spatial*_) or temporal (*f*_*temporal*_) data.

## 2 | MATERIAL & METHODS

### Time-series data

Time-series data were assembled from marine sediment cores containing planktonic foraminiferal assemblage data and independent sea-surface temperature reconstructions. To be included, cores had to satisfy five criteria: (1) they contained down-core records of planktonic foraminiferal assemblages and sea-surface temperature; (2) temperature reconstructions were based on proxies independent of foraminiferal assemblages, namely alkenones or Mg/Ca; (3) an age model was available, allowing assemblage and temperature records to be aligned through time; and (4) both assemblage and temperature series included depth or age information within the overlapping interval, allowing the two records to be merged. No cores meeting these criteria extended beyond 250,000 years, placing the resulting dataset well within the interval over which planktonic foraminiferal thermal niches are thought to have remained stable (Antell et al., 2021), with no speciation or extinction events (Lamyman et al., 2026). Candidate assemblage records were identified in the Triton database (Fenton et al., 2021) and in Strack et al. (2022), and the original assemblage data were downloaded from their primary repositories. Taxonomic synonymization of species names followed (Siccha and Kucera, 2017). For each core with assemblage data, we searched the PANGAEA data repository until the year 2025 for associated records of sea-surface temperature using the core name together with the terms, including “alkenone”, “alk*”, “UK37”, “Uk37”, “uk37”, “U*” + “37”, “Mg/Ca”, and “Mg” + “Ca”. To assign a temperature estimate to each assemblage sample, temperature records for each core were linearly interpolated to a regular depth resolution of 0.01 meter and matched to the depths of the planktonic foraminifera assemblage samples; for core 202-1240, for which depth information was unavailable for one of the series, ages were matched directly. The final time-series dataset comprises 12 sediment cores spanning 0.07 to 245 thousand years ago (ka BP; before 1950) and temperatures between 6.2 and 28.4 °C, with a minimum of 28 and a maximum of 652 samples per time series (Fig. 2b).

**F I G U R E 2.**
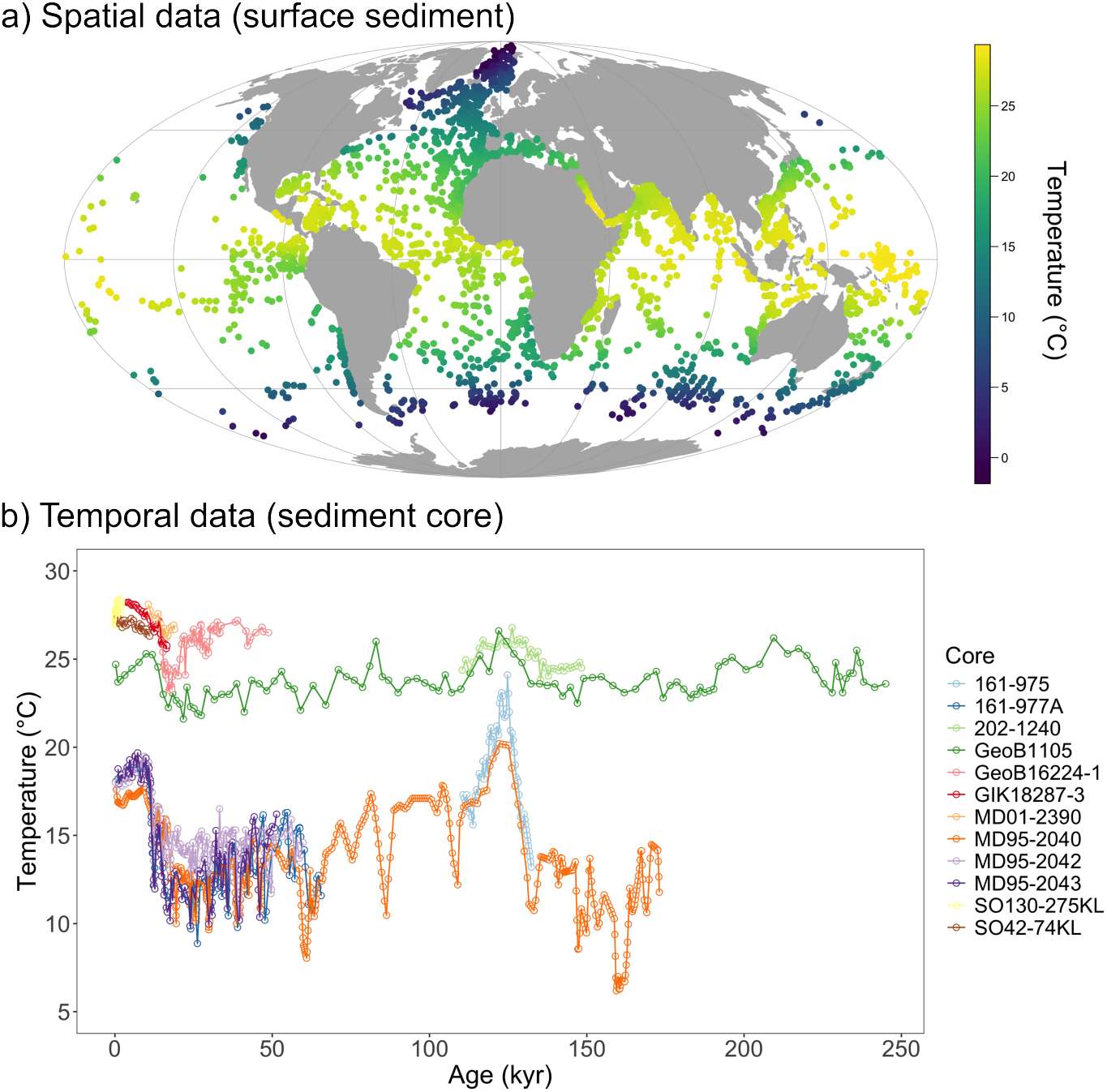
Overview of the spatial and temporal datasets. **(a)** Spatial dataset of surface-sediment sample locations (n = 3,515), coloured by mean annual sea-surface temperature (ERSSTv5, 1854–1899). **(b)** Site-specific sea-surface temperature reconstructions through time for the 12 sediment cores comprising the temporal dataset, coloured by core.

### Spatial data

We retrieved spatial assemblage data from the ForCenS database (Siccha and Kucera, 2017), a curated compilation of planktonic foraminiferal census counts from 4,205 marine surface-sediment samples. Following Rillo et al. (2022), we removed samples that shared identical coordinates (retaining one at random from each duplicate set) and samples that did not differentiate between species pairs, yielding a dataset of 41 planktonic foraminifera species that matches the taxonomy used for the fossil time series (see above). We additionally removed samples from waters deeper than 4000 m in the Pacific and Indian Oceans to reduce the effect of dissolution on assemblage composition (Rillo et al., 2022). We obtained associated temperature data from the Extended Reconstructed Sea Surface Temperature dataset (Huang et al., 2017) for the period 1854–1899 and estimated the mean annual temperature by averaging across all available years within this period. This period is the earliest available in ERSSTv5 and predates most anthropogenic warming, consistent with the multi-century-to-millennial averaging window of surface sediment deposition (Jonkers et al., 2019). The final spatial dataset consisted of 3,515 unique sites (Fig. 2a).

### Analyses

To ensure direct comparability of spatial and temporal data, all analyses were based on a stratified dataset in which spatial and temporal samples were balanced across the temperature gradient. Samples were grouped into 1 °C temperature bins; within each bin, the larger of the two datasets (spatial or temporal) was randomly subsampled to match the sample size of the smaller, and bins containing samples from only one dataset were dropped entirely, such that the two datasets spanned an identical temperature range with matched sample sizes per bin (Supp. Fig. S1). This stratification was performed once, before all subsequent train/test partitioning. All beta- and alpha-diversity analyses were performed on this stratified dataset. For both biodiversity dimensions, analyses were repeated across 100 train/test splits. In each replicate, temporal data were randomly partitioned into training and test subsets (50:50 split); spatial training data were then restricted to stratified spatial samples falling within the minimum-to-maximum temperature range of that replicate’s temporal training subset. Predictions were evaluated in three settings: time-for-time, in which temporal models were trained and tested on temporal data; space-for-time, in which spatial models were trained on spatial data and applied to temporal data; and space-for-space, in which spatial models were trained and tested within the spatial domain. Time-for-time and space-for-space performance served as within-domain benchmarks against which space-for-time performance was compared.

Predictive performance was quantified using pseudo-R^2^ (squared Pearson correlation between observed and predicted values on the response scale) and root mean square error (RMSE), calculated identically for beta- and alpha-diversity models on held-out test data. Because beta regression assumes a non-normal error structure, this pseudo-R^2^ does not carry the formal variance-explained interpretation of R^2^ in ordinary least squares, but provides a comparable, interpretable measure of predictive accuracy across model types. Fitted relationships were summarized across replicates by their median prediction and 95% interval. For supplementary observed-versus-predicted figures, a representative replicate was selected as the one with the space-for-time RMSE closest to the median across all replicates, to avoid selecting an unrepresentative best- or worst-case example.

### Beta diversity

To analyse compositional turnover across space and time, both the temporal and spatial assemblage datasets were converted into pairwise datasets in which each observation represents the dissimilarity between two samples. For each pair, we calculated abundance-based compositional dissimilarity using the Morisita-Horn index, implemented in the R package vegan (Oksanen et al., 2019), and the corresponding absolute difference in reconstructed sea-surface temperature, which we square-root transformed. This transformation improved model fit relative to untransformed temperature differences and, unlike a log transformation, permits temperature differences of zero. Pairwise compositional dissimilarity was modeled as a function of the square-root-transformed temperature difference using beta regression, implemented in the R package gamlss (Rigby & Stasinopoulos), with both the mean (logit link) and dispersion (log link) parameters modeled as functions of temperature. We used linear rather than smooth predictor terms, which allowed us to (i) directly compare temperature slopes between spatial and temporal models, and (ii) exchange slopes between models while holding intercepts fixed (see below); linear temperature-dissimilarity relationships are also supported at global scale by Rillo et al. (2022). Because beta regression requires responses to lie strictly within the open interval (0,1), exact dissimilarity values of 0 and 1 (0.04% of observations) were replaced by 0.001 and 0.999 before model fitting, following standard practice for boundary correction in beta regression (Smithson & Verkuilen, 2006). We conducted two complementary analyses: a global analysis, in which all time series were pooled and compared to the full spatial dataset (as in Blois et al., 2013), and a local analysis, in which each sediment core was analysed separately, with the corresponding spatial dataset restricted to the same ocean basin and temperature range as that core.

Pairwise temporal and spatial datasets were split into training and test sets by partitioning the original sample identities rather than the pairwise observations themselves, so that no sample contributed to both sets; pairwise observations were then retained only when both members of the pair belonged to the same partition. To reduce dependence among pairwise observations, training data were further thinned by randomly pairing each sample with another within the same partition, retaining only non-overlapping pairs, so that each sample contributed to only one pair per replicate. This train/test splitting and thinning procedure was repeated 100 times for both global and local analyses. In the global analysis, the spatial pairwise dataset was restricted to the range of temperature differences present in the temporal training data; in the local analysis, the same approach was applied separately to each sediment core. For each replicate, a temporal model was fitted to the temporal training data and a spatial model to the spatial training data. We then generated three types of temporal predictions: time-for-time, in which the temporal model predicted temporal test data; space-for-time, in which the spatial model predicted temporal test data; and space-slope-for-time, in which the temporal intercept was retained but the temporal slope was replaced by the spatial slope, isolating the transferability of the temperature-dissimilarity slope from the intercept offset between models. This prediction is most directly comparable to Blois et al. (2013), who reset model intercepts to zero before generating space-for-time predictions; we instead retained the fitted temporal intercept because forcing intercepts to zero distorts the model fit. The spatial model was also evaluated on spatial test data as a benchmark (space-for-space).

For the global analysis, replicate predictions were summarized along a common temperature-difference gradient. For local analyses, the same framework was applied separately to each sediment core; intercept and slope estimates were extracted from each replicate and model coefficient differences were calculated as spatial minus temporal. These differences were visualized as distributions across replicates, and tested against zero using Wilcoxon signed-rank tests. Because beta regression was fitted with a logit link for the mean, the model coefficients are interpreted on the link scale: intercepts represent the expected log-odds of dissimilarity at zero temperature difference, and slopes represent the change in log-odds of dissimilarity per unit increase in the square-root-transformed temperature difference.

Lastly, to assess whether our results were sensitive to temperature proxy choice, we repeated the local (single-core) space-for-time analysis for core GeoB16224-1. This core had more than one independently reconstructed temperature record (proxies: UK37, Mg/Ca, TEX86L; Crivellari et al. 2019). A TEX86H-based reconstruction was also available for this core but was excluded, as Crivellari et al. (2019) report it yields unrealistically high temperature values at this site due to contamination from terrigenous organic matter delivered by the Amazon River; TEX86L, which excludes the affected compound, was their preferred reconstruction. For each proxy, we recalculated the core’s temporal dissimilarity-temperature relationship and compared it to the same spatial subset used in the main local analysis (matched by ocean basin and temperature range), following the same procedure described above.

### Alpha diversity

Alpha diversity was quantified at the sample level using Hill numbers expressed as effective numbers of species (ENS), calculated as the exponential of Shannon entropy (q = 1) (Jost et al., 2011). We chose this metric because it incorporates both richness and evenness while remaining less dominated by rare taxa than species richness alone. The functional form of the alpha diversity-temperature relationship was selected by fitting linear, quadratic, and cubic polynomial models separately to the spatial and temporal datasets and comparing them using AIC, BIC, adjusted R^2^, and RMSE. Cubic models provided the best overall fit for ENS and were therefore used in all subsequent alpha-diversity analyses (Supp. Table S1). For each replicate, cubic-polynomial ordinary least-squares regression models of ENS on temperature were fitted separately to temporal and spatial training subsets, using a standard 50:50 random train/test split at the sample level. Predictions were then generated for the test subset of the temporal data (time-for-time and space-for-time) and the test subset of the spatial data (space-for-space); because cubic polynomial models do not yield a single slope parameter, the space-slope-for-time comparison used for beta diversity was not applicable here.

Regional alpha-diversity analyses followed the same framework as the global analysis; spatial and temporal samples were restricted to the Atlantic Ocean (pooling North Atlantic, South Atlantic, and Arctic groupings; Mediterranean samples were classified separately and excluded). This restriction reflects both ecological and methodological considerations: the temperature-diversity relationship in planktonic foraminifera is strongest in the Atlantic and weaker in the Pacific and Indian Ocean (Fenton et al., 2016), while the Mediterranean, Indian, and Pacific temporal datasets covered comparatively narrow temperature gradients, limiting their suitability for estimating nonlinear temperature-alpha-diversity relationships. Unlike beta diversity, where local analyses were performed per core, alpha-diversity regional analyses were conducted at the ocean-basin level, since estimating nonlinear alpha diversity-temperature relationships required broader thermal coverage than was available within most individual cores.

## 3 | RESULTS

### 3.1 | Spatial and temporal turnover relationships have similar slopes but different intercepts

At the global scale, temporal compositional dissimilarity increased with sea-surface temperature change, and spatial and temporal turnover relationships showed similar slopes but different intercepts (Fig. 3a). Space-for-time predictions were consistently shifted upward relative to the time-for-time relationship across the full temperature gradient, indicating higher baseline turnover in the spatial model. This intercept difference was significantly greater than zero across replicates (Wilcoxon signed-rank test: V = 5050, P < 0.0001), whereas the slope difference, though also statistically significant (V = 1443, P = 0.0002), had an interval that spanned zero (median = −0.04, 95% CI: −0.29 – 0.14; Fig. 4), consistent with the hybrid space-slope-for-time model closely tracking time-for-time (Fig. 3a) and its comparable RMSE (Fig. 3c). All models predicting turnover had modest pseudo-R^2^ values, indicating that although temperature change is a consistent correlate of turnover, it explains only part of temporal and spatial variation (Fig. 3b). Time-for-time had the lowest RMSE together with space-slope-for-time, while space-for-time performed worst (Fig. 3c). Observed-versus-predicted plots further showed that these models strongly compressed the range of predicted dissimilarities relative to the observed data (Supp. Fig. S2).

**F I G U R E 3.**
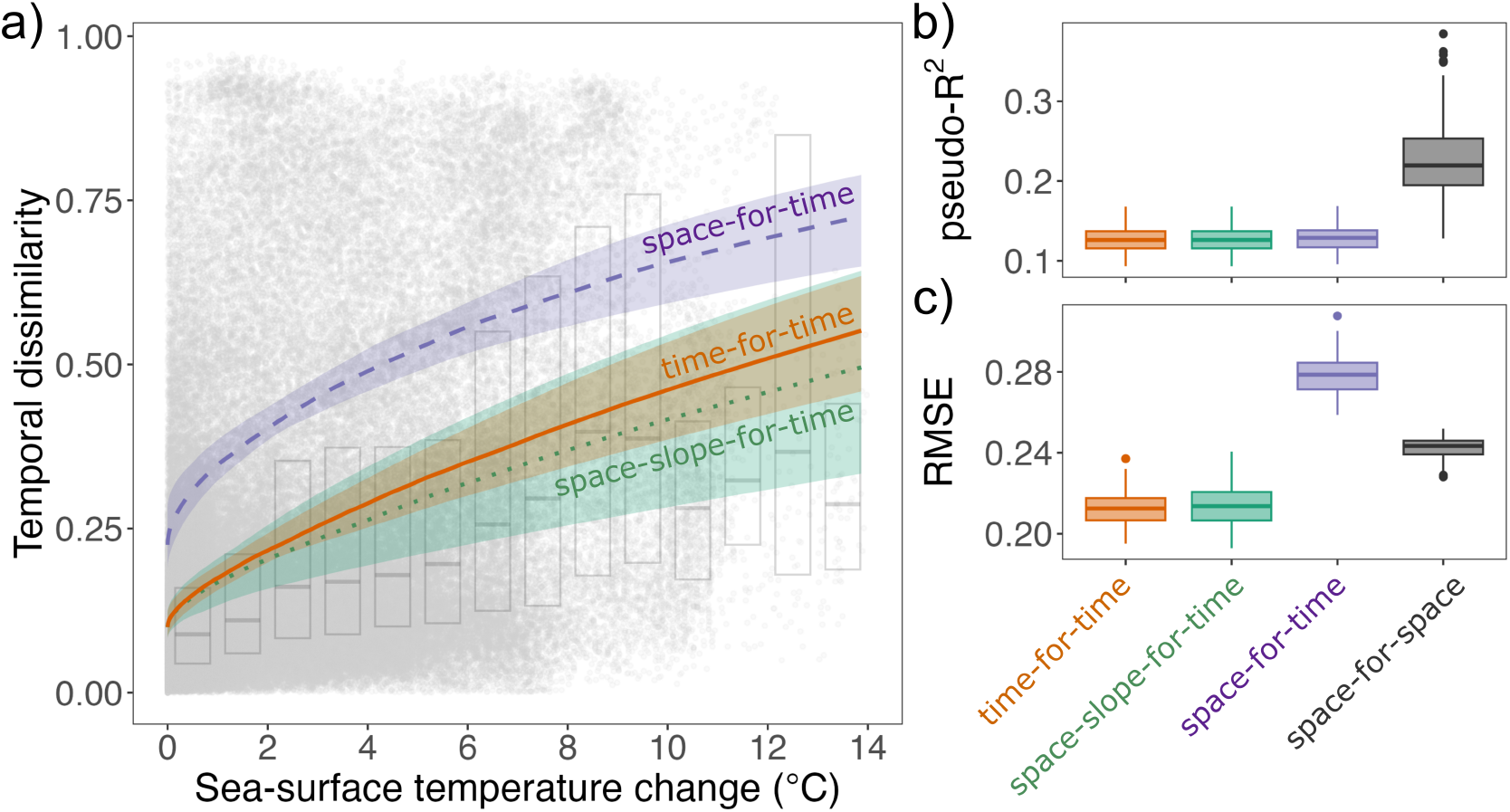
Global comparison of temporal and spatial turnover-temperature relationships. **(a)** Temporal pairwise compositional dissimilarity is shown as a function of sea-surface temperature change, with median fitted relationships across 100 replicate train/test splits for the temporal model (time-for-time), the spatial model applied to temporal data (space-for-time), and the hybrid model retaining the temporal intercept but substituting the spatial slope (space-slope-for-time). Shaded ribbons show the 95% interval across replicate fits. Faint background boxplots summarize the distribution of observed temporal dissimilarity within paleo-temperature-change bins. Side panels show the distribution across replicates of predictive performance, quantified as **(b)** pseudo-R^2^ and **(c)** root mean squared error (RMSE), for time-for-time, space-for-time, space-slope-for-time, and the spatial model evaluated on spatial data (space-for-space).

**F I G U R E 4.**
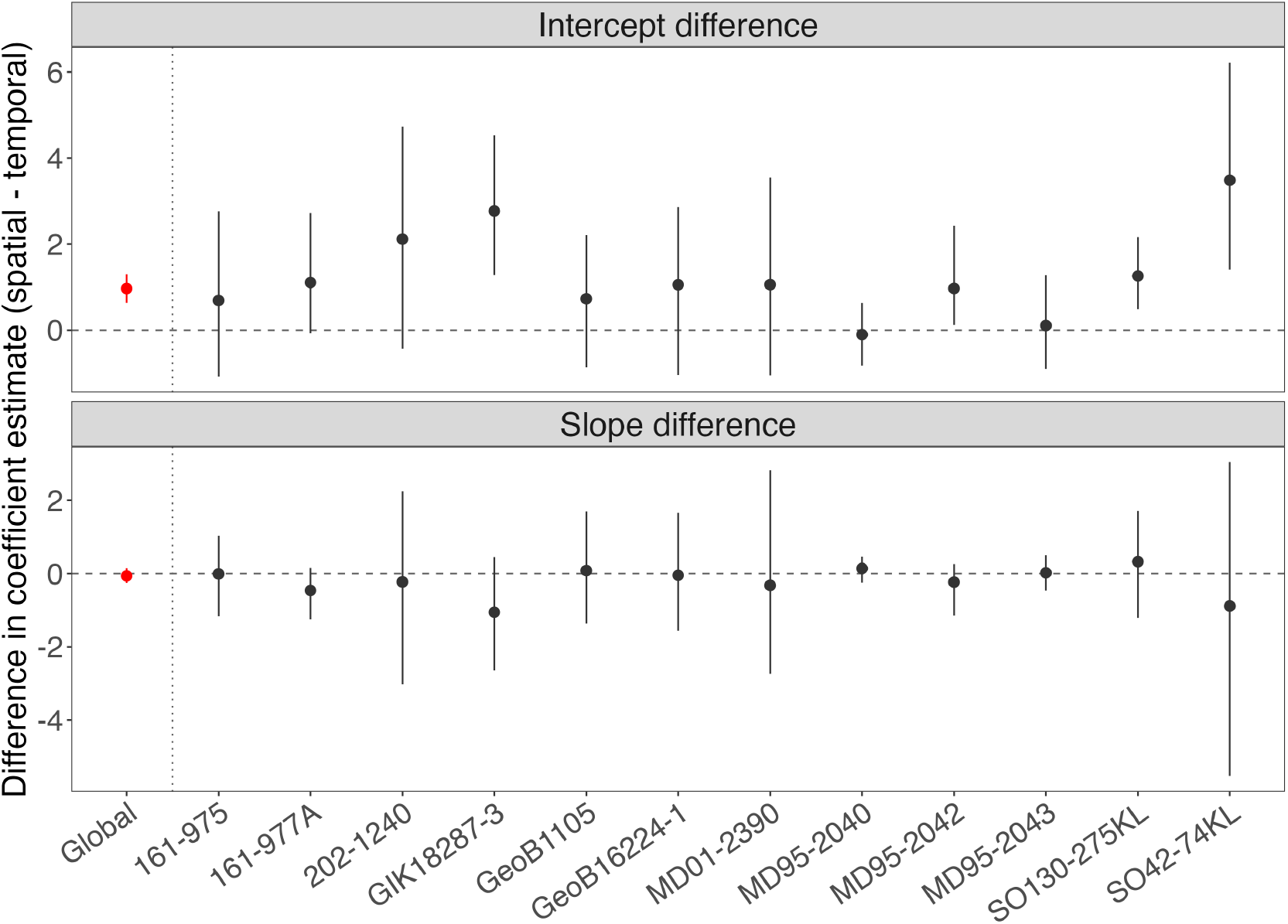
Global and local spatial-temporal differences in model coefficients across time series. Globally (i.e., all cores together) or for each core, differences in coefficient estimates between the spatial and temporal models are shown for the intercept and slope, calculated as spatial minus temporal. Points are median differences across replicate fits, and vertical bars indicate the 95% interval across replicates. The dashed horizontal line marks zero, so positive values indicate higher coefficient estimates in the spatial model than in the temporal model. The dotted vertical line separates the global analyses from the local (core-specific) analyses. Globally and across cores, intercept differences were significantly greater than zero, whereas slope differences were not significantly different from zero.

### 3.2 | Local spatial restriction reduces but does not eliminate baseline turnover mismatch

Because the higher spatial intercept could arise from the broad geographic extent of the global spatial dataset, we next repeated the comparison using spatial data restricted to the ocean basin and temperature variability of each fossil time series. This local analysis narrowed the intercept mismatch between spatial and temporal turnover relationships, but did not eliminate it (Fig. 4). Across cores, local spatial-temporal differences in slope were generally small and centered near zero, whereas differences in intercept remained predominantly positive. Intercept differences were significantly greater than zero across cores (Wilcoxon signed-rank test: V = 78, P = 0.0005), whereas slope differences were not significantly different from zero (V = 17, P = 0.092; Fig. 4). Core-specific fitted relationships and predictive performance showed that this overall pattern was broadly consistent across time series despite variation in local predictability (Supp. Figs. S3 and S4). Thus, restricting the spatial comparison locally reduced the mismatch between space and time, but local spatial models still tended to predict higher baseline turnover than temporal models.

### 3.3 | Alpha diversity relationships are more transferable than turnover relationships

The effective number of species (ENS) increased nonlinearly with sea-surface temperature in both the spatial and temporal datasets, and model comparisons supported the use of cubic models for subsequent analyses (Supp. Table S1).

In the global comparison, the spatial and temporal alpha diversity-temperature relationships were broadly similar in overall form, although predicted ENS differed by up to 2.4 species-equivalents between spatial and temporal models at the warm and cool ends of the gradient (Fig. 5a). This mismatch at the higher and lower ends of the temperature gradient may indicate temperature-proxy biases (Tierney and Tingley, 2018) and/or time-series-specific bias, as the warm and cool ends of the gradient are represented predominantly by a single core each (Fig. 2b). The fitted relationships were also highly consistent across replicate train/test splits, as indicated by the narrow 95% intervals around the median curves (Fig. 5a). Unlike compositional turnover, alpha diversity did not show a strong offset between spatial and temporal relationships, and spatial models therefore did not systematically overestimate the temporal response across the full gradient. Across replicates, time-for-time predictions yielded higher R^2^ and lower RMSE than space-for-time predictions, while space-for-space performed best within the spatial domain (Fig. 5b,c; Supp. Fig. S5). Restricting the alpha-diversity analysis to the Atlantic increased predictive performance relative to the global analysis, but did not eliminate differences in fitted spatial and temporal alpha-temperature relationships (Supp. Fig. S6). Spatial models thus recovered the temporal alpha-diversity relationship more closely than they did for compositional turnover, indicating greater transferability from space to time for alpha diversity than for beta diversity, where spatial turnover models matched temporal slopes but overpredicted baseline turnover.

**F I G U R E 5.**
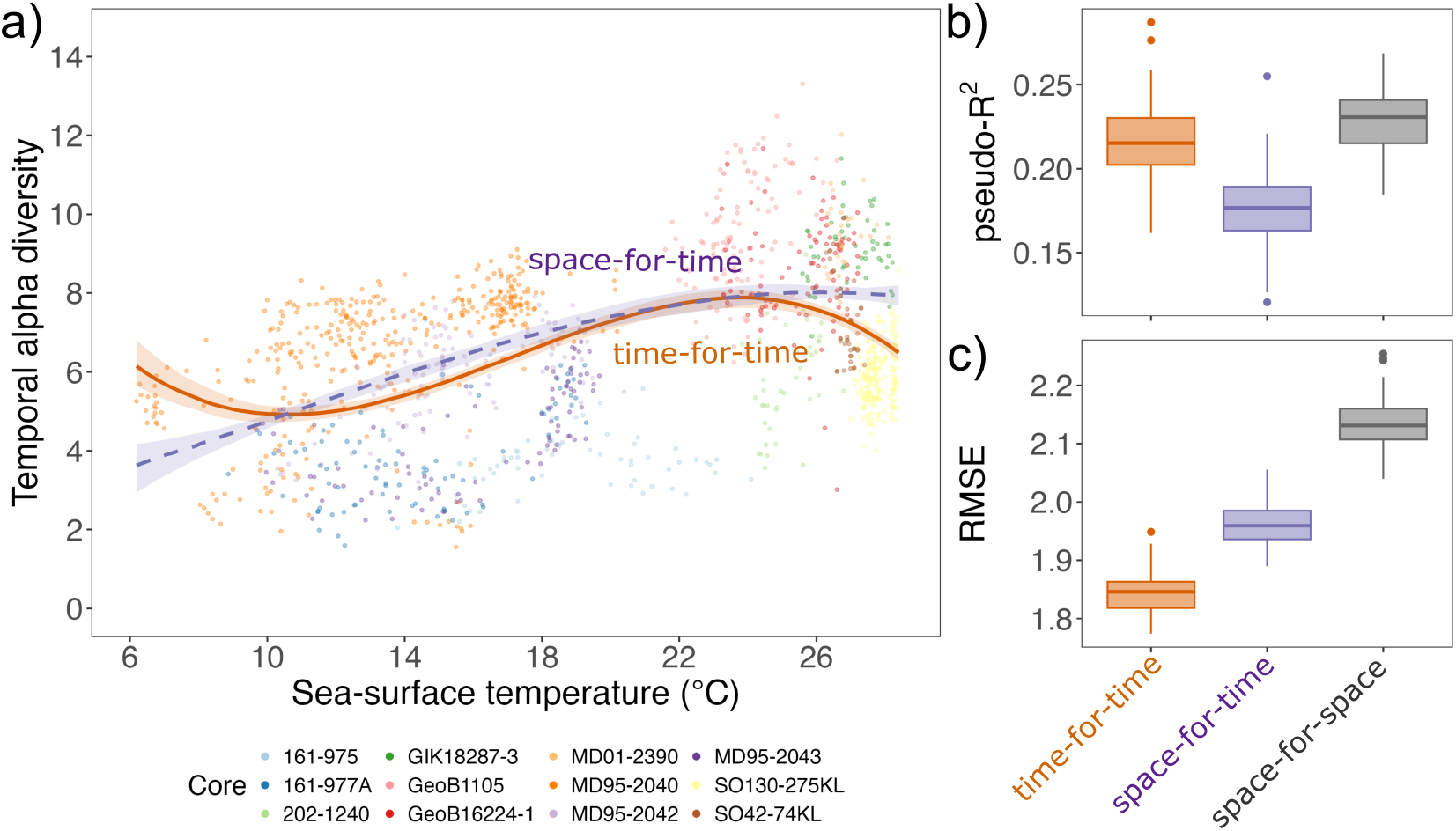
Global comparison of spatial and temporal alpha-temperature relationships. **(a)** Temporal alpha diversity (ENS, q=1) is plotted against sea-surface temperature. Lines show the median fitted relationships across 100 replicate train/test splits for the temporal model (time-for-time) and the spatial model applied to temporal data (space-for-time) fitted as cubic polynomial ordinary least-squares regressions; ribbons indicate the 95% interval across replicate fits. Side panels show predictive performance across replicates, expressed as **(b)** pseudo-R^2^ and **(c)** root mean squared error (RMSE), for time-for-time, space-for-time, and the spatial model evaluated within the spatial domain (space-for-space).

## 4 | DISCUSSION

Despite previous empirical validations, the predictive power of space-for-time substitution remains poorly constrained. This limitation is concerning, given the urgency of anticipating biodiversity responses to anthropogenic climate change and the extent to which such forecasts rely on this approach. Here, we show that space-for-time substitution can recover broad biodiversity-temperature relationships through time, but that its transferability is incomplete and depends on which component of biodiversity change is being predicted. For beta diversity, spatial and temporal turnover relationships showed similar slopes, but spatial models consistently predicted higher baseline turnover than temporal models (Figs. 3, 4). For alpha diversity, spatial models recovered the general nonlinear form of the temporal biodiversity-temperature relationship more closely than for compositional turnover (Fig. 5).

### 4.1 | Spatial and temporal turnover differ mainly in baseline turnover

For beta diversity, spatial and temporal turnover relationships showed similar slopes but different intercepts in both the global (Fig. 3) and local (Fig. 4) analyses. Spatial models therefore recovered the temperature sensitivity of temporal turnover, but overpredicted its baseline level. This result differs from Blois et al. (2013), who fixed the intercept at zero, thereby guaranteeing that zero climate difference gives zero predicted turnover and potentially increasing the apparent fit between spatial and temporal models. By estimating the intercept freely, our models revealed that the main mismatch is not how strongly turnover changes with temperature, but how much turnover is already expected when temperature difference is zero.

Spatial climate-biotic relationships are often interpreted as reflecting the outcome of ecological processes operating over longer timescales, including colonization, extinction, and prolonged responses to environmental change, whereas temporal observations over shorter intervals may be more strongly shaped by fast processes such as demographic responses, short-term environmental variation, and transient dynamics (Levin 1993; Adler et al. 2020). Although temporal turnover is obviously related to spatial dissimilarity at shorter time scales, as new species can only be observed in time if they are in spatial proximity (Hodapp et al ELE 2018), this timescale difference has often been invoked to explain low transferability from space to time (Damgaard 2019; Evans et al. 2025; Lovell et al. 2023; Perret et al. 2024). Our sediment core assemblages, however, integrate community change over hundreds to thousands of years (except core SO130-275KL), a timescale over which slower processes such as colonization, local extinction, and prolonged responses to environmental change are more likely to be expressed. This longer temporal scale can explain why spatial and temporal turnover models showed similar slopes (Fig. 3). However, this convergence did not extend to the intercept, as spatial models consistently predicted higher baseline turnover than temporal models (Fig. 3), even when the spatial comparison was restricted to local areas (Fig. 4). This offset suggests that temperature alone does not capture all of the background differences among sites that contribute to spatial turnover, including spatial sampling by different people as opposed to the sediment core data. Recent anthropogenic change could also contribute to this offset; however, the spatial assemblages in our study derive from pre-industrial surface sediments (Jonkers et al., 2019), making this cause unlikely. Our results indicate that compatible timescales may improve the transferability of biodiversity-temperature slopes, but they do not guarantee equivalence in the absolute magnitude of biodiversity change.

### 4.2 | Temperature is an important but incomplete predictor of biodiversity change

Temperature is repeatedly identified as the main predictor of planktonic foraminiferal diversity in space (Rutherford et al., 1999; Fenton et al., 2016; Morey et al., 2005; Rillo et al., 2022); however, these relationships are based on correlative analyses. We found a broad agreement in slope direction between spatial and temporal beta and alpha diversity (Figs. 3 and 5), which is consistent with temperature having a causal role in shaping planktonic foraminifera biodiversity change (Lovell et al., 2023) and with the stability of species’ thermal niches over time (Antell et al., 2021).

Despite this agreement in slope, the low R^2^ values for all turnover models (Fig. 3b, Supp. Fig. S2) show that temperature remains an incomplete predictor of species composition through space and time. Other abiotic variables, biotic interactions, biogeographic structure, and interactions among predictors may all contribute to turnover (Rillo et al., 2022; Strack et al., 2022; Jonkers et al., 2023; Ying et al., 2024) and weaken the predictive power of a temperature-only model. Moreover, when modelling turnover, each pairwise comparison is reduced to a single temperature difference, creating a many-to-one problem: the same temperature difference can arise from very different absolute temperature combinations. As a result, different assemblage pairs are forced onto the same predictor value (i.e., same temperature difference; Supp. Fig. S2), even though they may differ markedly in composition depending on where they occur along the full temperature gradient.

Lastly, the limited ability of temperature to predict biodiversity change seen in our turnover models may seem surprising given the long use of planktonic foraminifera in paleoenvironmental reconstruction (Kucera, 2007). Transfer functions and modern analogue approaches solve a different inferential problem: they use assemblage composition as a predictor of past environmental conditions, whereas our analyses use temperature change as a predictor of biodiversity change. However, such paleoenvironmental approaches typically ignore the space-for-time substitution assumption (Juggins, 2013; Blonder et al., 2017), demonstrating the importance of empirically validating this assumption.

### 4.3 | Alpha diversity is more transferable than turnover

The loss of thermal information in the turnover models discussed above may also help explain why alpha diversity was more transferable than beta diversity. Alpha models retain the absolute temperature of each sample, whereas turnover is modelled from a single temperature-difference value for each pair of samples. As a result, alpha models preserve more of the thermal context in which biodiversity variation occurs. At the same time, alpha diversity summarizes biodiversity more coarsely than beta diversity because it does not retain species composition, only the effective number of species (Hillebrand et al., 2018). Thus, alpha diversity may be easier to predict across space and time because it reduces biodiversity to a single summary value, whereas compositional turnover directly quantifies changes in species composition. Our results suggest that transferability may depend both on how much climatic information is included in the predictor and on how much compositional information is retained in the response. This interpretation is broadly consistent with previous work showing stronger mismatches between spatial and temporal models at finer biological resolution and over shorter timescales, such as the species- and population-level responses examined by Perret et al. (2024). More generally, short-term dynamics may be shaped more strongly by fast processes and transient responses, whereas broader community patterns integrate slower processes operating over longer timescales (Adler et al., 2020). Community responses may therefore be more predictable both because individual species’ responses are heterogeneous, but their aggregation produces more predictable community patterns (Srivastava et al., 2021; Ferrier et al., 2007) and because community-level patterns are more likely to reflect the slower processes captured by spatial gradients.

### 4.4 | Robustness to temperature proxy choice

Sea-surface temperature estimates from different proxies carry different sources of uncertainty that may influence our results. Eleven of our 12 cores are based on alkenone (UK’37) reconstructions, which tend to underestimate warming at high temperatures and cooling at low temperatures (Tierney and Tingley, 2018). This bias may explain the space-time mismatch and the sigmoidal shape of the alpha diversity curve at higher temperatures (Fig. 5).

To test whether our conclusions were sensitive to proxy choice, we repeated the local space-for-time analysis for core GeoB16224-1, which contained independently reconstructed temperature records from three proxies (UK’37, Mg/Ca, TEX86L; Crivellari et al. 2019; see Methods). Across all three proxies, spatial models consistently predicted higher dissimilarity than temporal models (Wilcoxon signed-rank test: UK’37 p < 0.001, MgCa p < 0.001, TEX86L p = 0.033; Supp. Fig. S7), although the size of this offset was itself proxy-dependent (Kruskal-Wallis test, 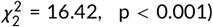 being largest for UK’37 – the proxy used in our main analysis – which differed significantly from both alternative proxies (pairwise Wilcoxon, both p < 0.002 vs. UK’37), while MgCa and TEX86L did not differ from one another. In contrast, the similarity we report between spatial and temporal slopes for this core held only for UK’37 (p = 0.22); the spatial and temporal slopes differed significantly for both alternative proxies (MgCa p = 0.002, TEX86L p < 0.001), consistent with the slope difference varying significantly among proxies (Kruskal-Wallis, 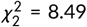, p = 0.014). Thus, for this core, the intercept difference between spatial and temporal models was robust to proxy choice, while the slope difference was not.

We further tested how alkenone-based proxy recontruction could bias our main, 12-core result by splitting cores into a “cool” group (median temporal temperature ≤ 24°C; n = 6, five alkenone-based and one Mg/Ca-based core, GeoB1105) and a “warm” group (median temperature > 24°C; n = 6, all alkenone-based) (Supp. Table S2); and tested whether the intercept and slope effects where different among these two groups. The intercept difference was significant in the warm group (Wilcoxon signed-rank test, p = 0.031) but not in the cool group (p = 0.063), where the effect was also smaller in magnitude (median −0.10 – 1.11 vs. 1.06 – 3.48 across cores). The slope difference remained non-significant in both temperature groups (warm, median −1.06 – 0.33, p = 0.219; cool, median −0.46 – 0.14, p = 0.844), consistent with our full 12-core result. Although six cores per group reduce statistical power, these results suggest that, when temperature estimates are unbiased, the slope and magnitude of the biodiversity-temperature relationship are similar – making space-for-time more transferable.

### 4.5 | Implications for forecasting biodiversity change

Our results suggest that, over the long timescales represented here, spatial gradients can recover the slope of temporal marine biodiversity-temperature relationships, but not their intercept. In practice, spatial models may be informative about the direction and relative change of biodiversity, particularly when the forecast horizon is long enough for slower ecological processes to occur (Adler et al., 2020; Evans et al., 2025). A key implication of our finding that spatial and temporal slopes are similar is that spatial models can reliably predict the direction and relative rate of biodiversity change through time. Forecasts anchored to the observed present-day state (intercept) and using spatial gradients to estimate relative changes as anomalies rather than absolute values carry substantially lower uncertainty, making this a more useful approach for biodiversity forecasting under climate change.

## Acknowledgements

We thank Tonke Strack and Iván Hernández-Almeida for helping with finding suitable sediment core data, and Rodrigo Costa Portilho-Ramos for sharing unpublished data. MCR was funded by the German Research Foundation (DFG) Cluster of Excellence ‘The Ocean Floor—Earth’s Uncharted Interface’ (EXC 2077, grant no. 390741603). LJ was supported by the German Federal Ministry of Research, Technology and Space (BMFTR) as a Research for Sustainability initiative (FONA) through the PALMOD project (grant no. 01LP2308A).

## Authors’ contribution

MCR conceived the study, compiled the sediment core dataset, and performed analyses. LM contributed to data curation, analyses, method development, and interpretation of results. LJ contributed to data curation and interpretation of results. JM contributed to analyses and method development. HH contributed to method development, interpretation of results, and secured funding. MCR wrote the first draft of the manuscript, and all authors read and edited the manuscript.

## Data availability

All code supporting this manuscript is archived on Zenodo (https://doi.org/10.5281/zenodo.21977364). This study compiled data from multiple publicly available sources. Complete information on all datasets, including DOIs, database sources, and data descriptions, is provided in Supplementary Table S3.

