## Supplementary Material for "Space-for-time substitution reveals partial transferability of marine biodiversity-temperature relationships"

465 # Supplementary Material

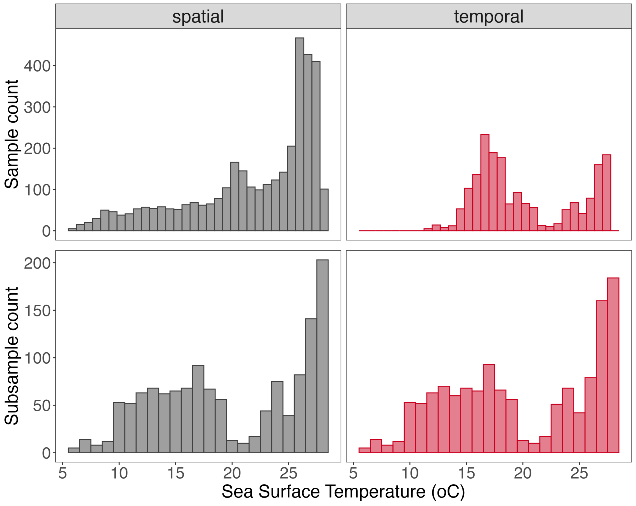


**FIGURE S1 Stratification of the spatial and temporal datasets to a common sea-surface temperature distribution.** Top row: sample counts per 1 °C sea-surface temperature bin in the original spatial (grey) and temporal (red) datasets. Bottom row: sample counts per bin after stratification, in which the larger dataset was randomly subsampled within each bin to match the sample size of the smaller, yielding spatial and temporal datasets with matched sample sizes across an identical temperature range.

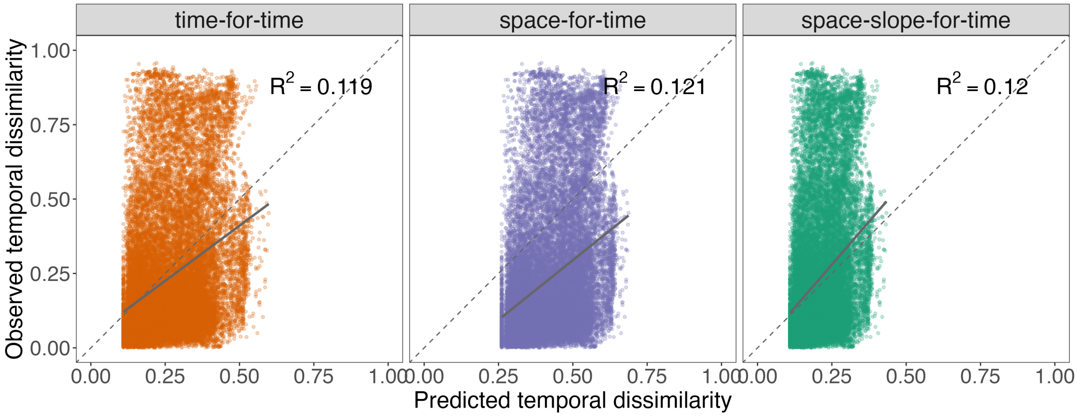


**FIGURE S2 Predicted versus observed temporal dissimilarity for the global beta-diversity analysis.** Predicted temporal dissimilarity is shown against observed temporal dissimilarity for the time-for-time, space-for-time, and space-slope-for-time models, using the representative replicate whose space-for-time RMSE was closest to the median across all replicates. Solid lines show least-squares fits, and dashed lines indicate the 1:1 expectation. In all three cases, predictions occupy a narrower range than the observed values, indicating compression of predicted dissimilarity relative to the full spread of temporal turnover.

**TABLE S1** Model selection criteria for linear, quadratic, and cubic polynomial ordinary least-squares regressions describing the relationship between alpha diversity (ENS1) and sea-surface temperature in the spatial and temporal datasets. ΔAIC and ΔBIC are shown relative to the best-supported model within each dataset; lower ΔAIC, ΔBIC, and RMSE values and higher adjusted $R^2$ indicate better model fit.

| Data | Model | ΔAIC | ΔBIC | Adjusted $R^2$ | RMSE |
|---|---|---|---|---|---|
| spatial | Linear | 1621.086 | 1607.463 | 0.523 | 3.823 |
| spatial | Quadratic | 518.453 | 511.642 | 0.596 | 3.521 |
| spatial | Cubic | 0 | 0 | 0.626 | 3.387 |
| temporal | Linear | 362.109 | 350.064 | 0.270 | 3.098 |
| temporal | Quadratic | 276.201 | 270.678 | 0.303 | 3.026 |
| temporal | Cubic | 0 | 0 | 0.400 | 2.806 |

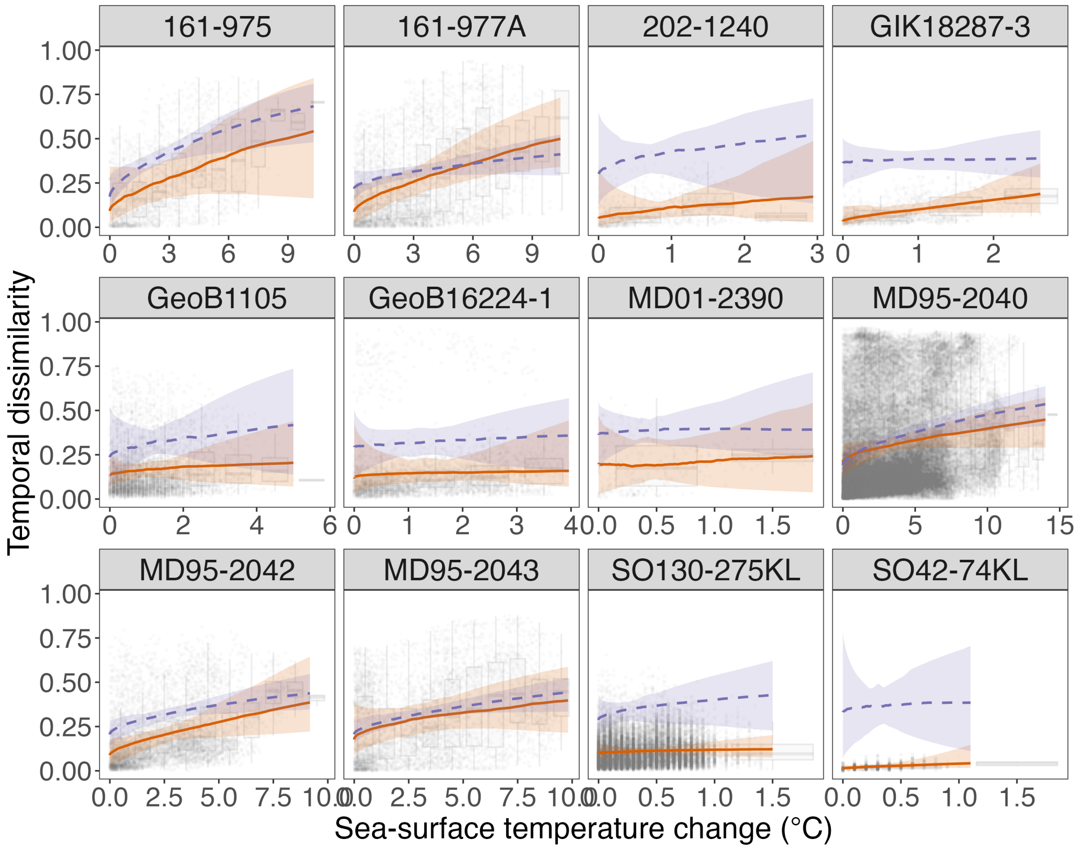


**FIGURE S3 Core-specific local spatial and temporal turnover relationships.** Temporal pairwise compositional dissimilarity is plotted against sea-surface temperature change separately for each sediment core. Lines show the median fitted relationships across replicate train/test splits for the temporal model (time-for-time: orange solid line) and the local spatial model applied to temporal data (space-for-time: purple, dashed line); ribbons indicate the 95% interval across replicate fits. Faint background boxplots and points summarize the distribution of temporal dissimilarity within each core.

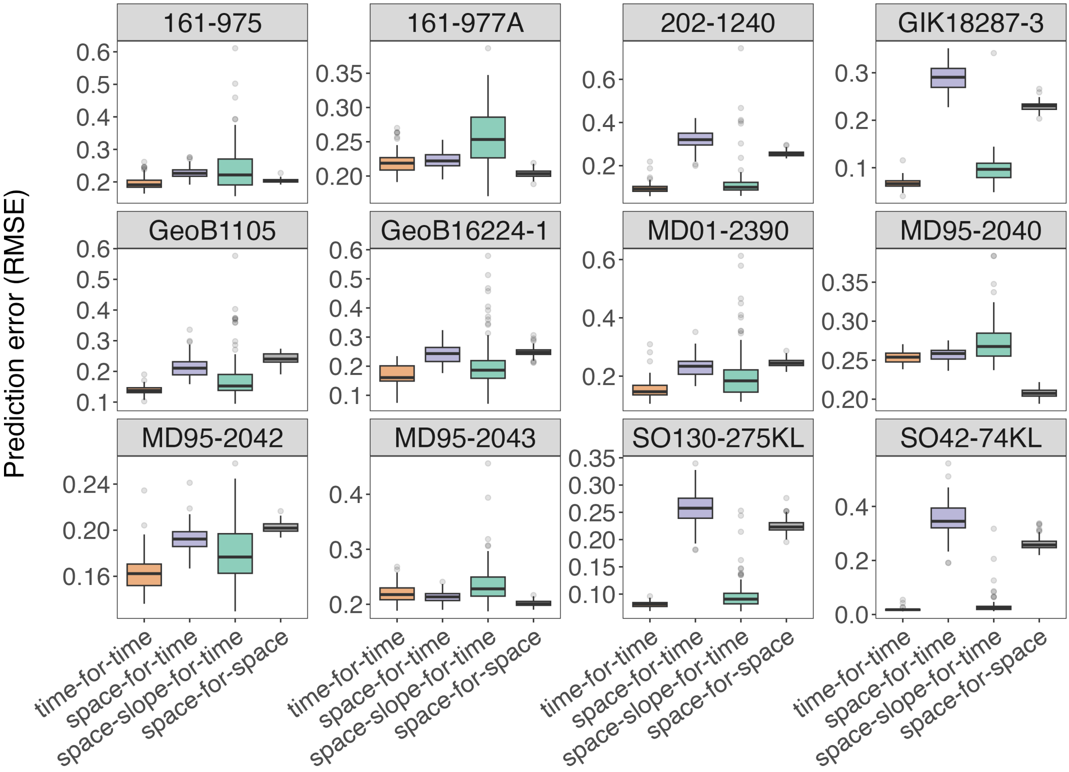


**FIGURE S4 Core-specific predictive performance of local spatial and temporal turnover models.** Boxplots show RMSE across replicate train/test splits for each sediment core for the temporal model applied to temporal data (time-for-time), the local spatial model applied to temporal data (space-for-time), the hybrid model retaining the temporal intercept but substituting the local spatial slope (space-slope-for-time), and the local spatial model evaluated within the spatial domain (space-for-space). Lower values indicate better predictive performance.

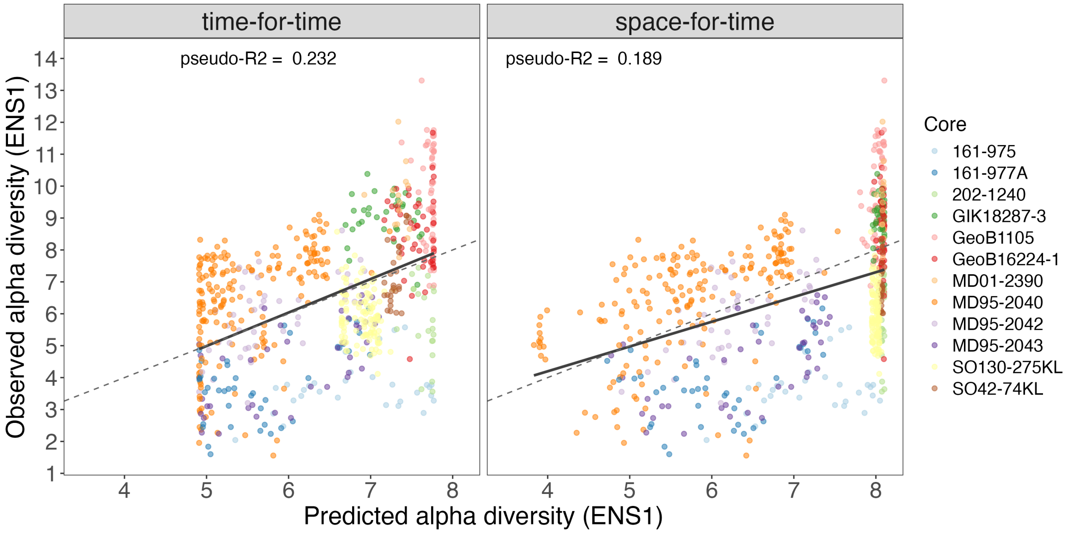


**FIGURE S5 Observed versus predicted alpha diversity in the global analysis.** Panels show observed and predicted ENS values for a representative replicate close to the median predictive performance across train/test splits, for the time-for-time and space-for-time models. Points are coloured by sediment core. Solid lines show ordinary least-squares fits and dashed lines indicate the 1:1 relationship. Pseudo-$R^2$ values are shown for each panel.

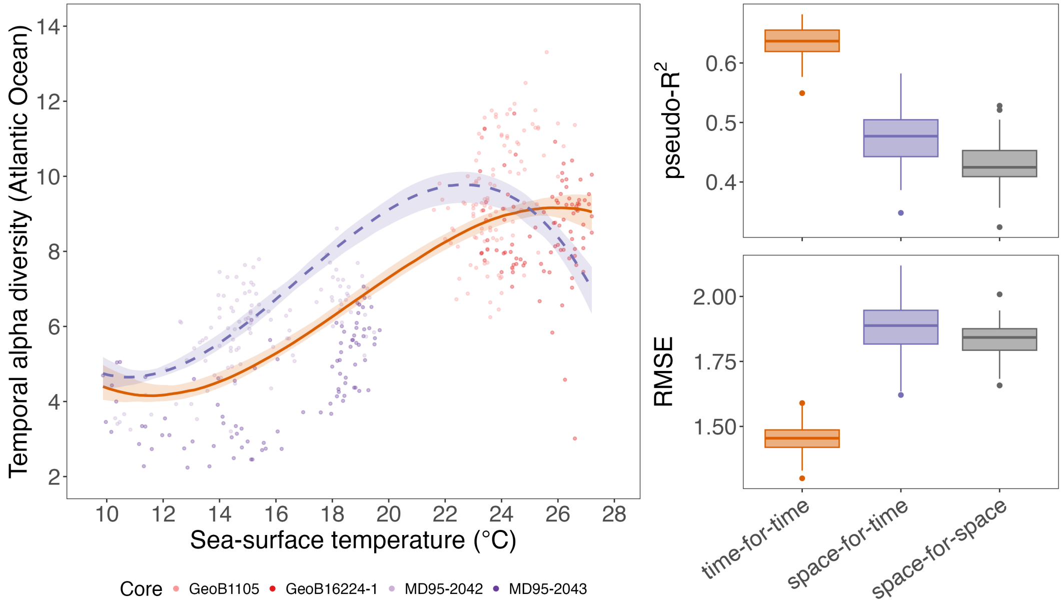


**FIGURE S6 Atlantic-only comparison of spatial and temporal alpha diversity-temperature relationships.** (a) Atlantic alpha diversity (Effective Number of Species q = 1) is plotted against sea-surface temperature for samples assigned to the Atlantic basin, excluding the Mediterranean. Lines show the median fitted relationships across replicate train/test splits for the temporal model (time-for-time) and the spatial model applied to temporal data (space-for-time); ribbons indicate the 95% interval across replicate fits. Side panels show predictive performance across replicates, expressed as (b) pseudo-$R^2$ and (c) Root Mean Squared Error (RMSE), for time-for-time, space-for-time, and the spatial model evaluated within the spatial domain (space-for-space).

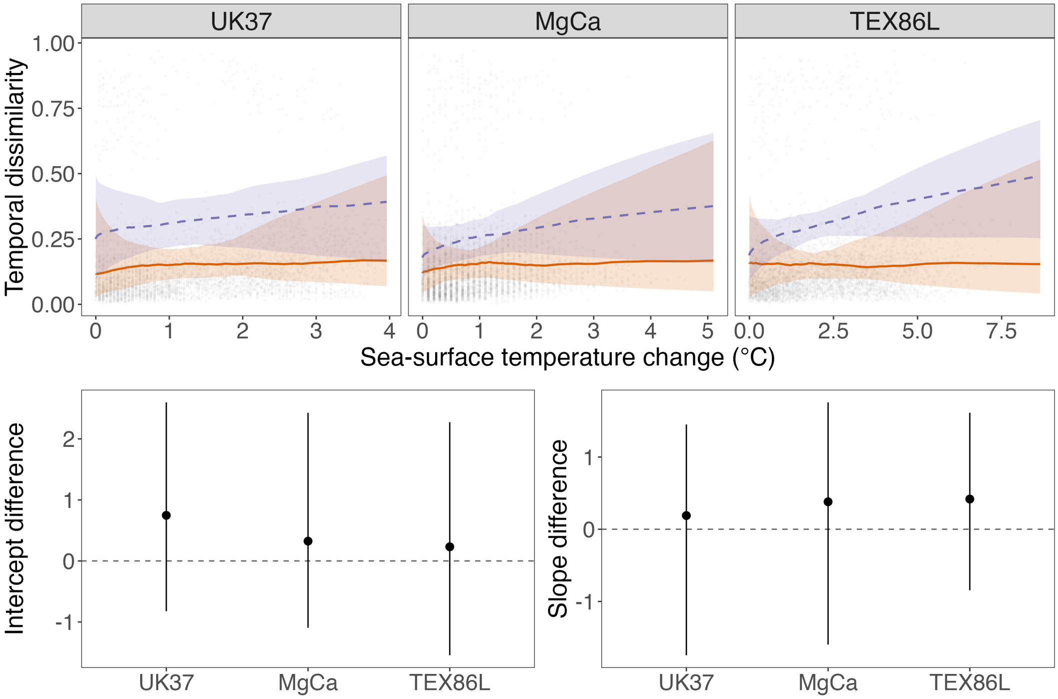


**FIGURE S7 Sensitivity of the local space-for-time comparison for core GeoB16224-1 to temperature proxy choice.** Top: temporal (orange, solid) and spatial (purple, dashed) dissimilarity–SST relationships fitted separately for each of the three proxies (UK'37, Mg/Ca, TEX86L; median and 95% CI across 100 replicate fits); grey dots show the underlying temporal pairwise dissimilarities. Bottom: median difference (spatial − temporal) in model intercept and slope for each proxy, with 95% CI; dashed line indicates no difference. The spatial model predicted significantly higher dissimilarity (intercept) than the temporal model for all three proxies (Wilcoxon signed-rank test: UK'37 $p < 0.001$, MgCa $p < 0.001$, TEX86L $p = 0.033$), though the size of this offset varied significantly among proxies (Kruskal-Wallis test, $\chi^2_2 = 16.42$, $p < 0.001$): UK'37 – the proxy used in the main analysis – showed a larger offset than both other proxies (pairwise Wilcoxon, BH-adjusted, both $p < 0.002$ vs. UK'37), while MgCa and TEX86L did not differ from one another. The spatial-temporal slope difference was significant for MgCa and TEX86L (both $p < 0.003$) but not for UK'37 ($p = 0.22$), consistent with the slope difference varying significantly among proxies (Kruskal-Wallis, $\chi^2_2 = 8.49$, $p = 0.014$).

**TABLE S2** Per-core spatial–temporal intercept and slope differences (local beta-diversity models) grouped by sea-surface temperature (SST) proxy and SST regime. “Cool” cores have median temporal temperature ≤ 24°C; “warm” cores have median temporal temperature > 24°C. Values are the median difference (spatial − temporal) across 100 replicate fits, with 95% CI in brackets. These values can be visualized in Fig. 4 of the main text.

| Core | Proxy | Median SST (°C) | Intercept diff. [95% CI] | Slope diff. [95% CI] |
|---|---|---|---|---|
| **Cool regime (median SST ≤ 24°C)** | | | | |
| MD95-2040 | Alkenone (UK’37) | 13.2 | −0.10 [−0.83, 0.63] | 0.14 [−0.25, 0.46] |
| 161-977A | Alkenone (UK’37) | 13.7 | 1.11 [−0.07, 2.73] | −0.46 [−1.25, 0.16] |
| MD95-2042 | Alkenone (UK’37) | 15.1 | 0.97 [0.12, 2.42] | −0.23 [−1.15, 0.26] |
| MD95-2043 | Alkenone (UK’37) | 17.3 | 0.11 [−0.90, 1.28] | 0.02 [−0.46, 0.50] |
| 161-975 | Alkenone (UK’37) | 18.6 | 0.69 [−1.08, 2.77] | −0.01 [−1.17, 1.03] |
| GeoB1105 | Mg/Ca | 23.6 | 0.73 [−0.86, 2.21] | 0.08 [−1.36, 1.70] |
| **Warm regime (median SST > 24°C)** | | | | |
| 202-1240 | Alkenone (UK’37) | 25.0 | 2.12 [−0.43, 4.73] | −0.23 [−3.02, 2.25] |
| GeoB16224-1 | Alkenone (UK’37) | 26.0 | 1.06 [−1.04, 2.86] | −0.05 [−1.56, 1.66] |
| MD01-2390 | Alkenone (UK’37) | 26.7 | 1.06 [−1.05, 3.55] | −0.32 [−2.74, 2.82] |
| SO42-74KL | Alkenone (UK’37) | 26.9 | 3.48 [1.41, 6.22] | −0.88 [−5.53, 3.04] |
| GIK18287-3 | Alkenone (UK’37) | 26.9 | 2.77 [1.28, 4.53] | −1.06 [−2.64, 0.45] |
| SO130-275KL | Alkenone (UK’37) | 27.7 | 1.26 [0.49, 2.16] | 0.33 [−1.21, 1.71] |

Group-level Wilcoxon signed-rank test (median difference vs. 0): **Cool group** ($n$ = 6) intercept $V$ = 20, $p$ = 0.063; slope $V$ = 9, $p$ = 0.844. **Warm group** ($n$ = 6) intercept $V$ = 21, $p$ = 0.031; slope $V$ = 4, $p$ = 0.219.

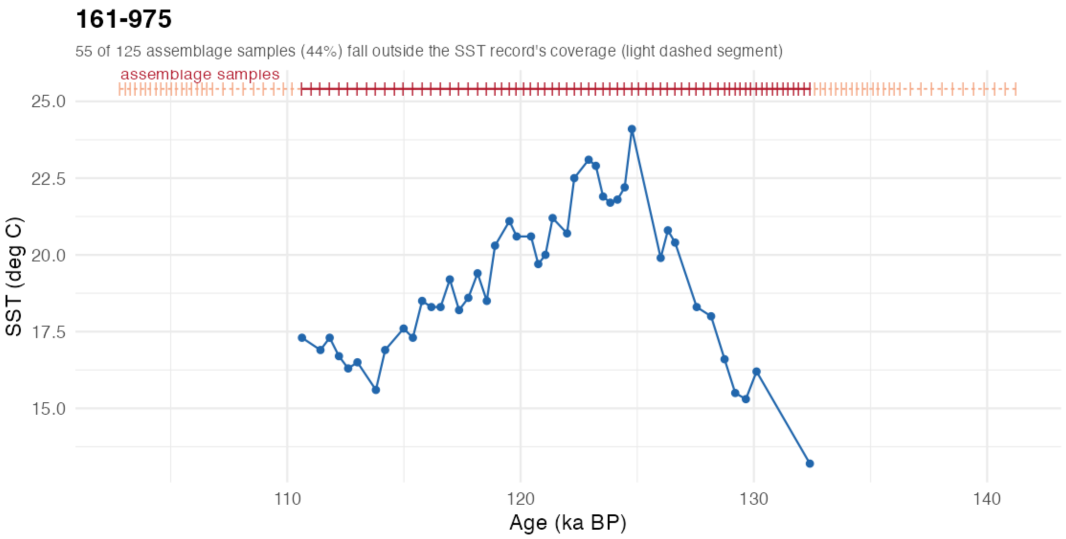


**FIGURE S8** Sea-surface temperature (SST) record coverage relative to planktonic foraminifera assemblage data, for each of the 12 sediment cores (alphabetic order). For each core, the SST record (line and points) is shown with a marker line above indicating the ages of all raw assemblage samples: solid dark red where assemblage ages fall within the SST record's coverage, and light dashed orange where they extend beyond it (samples that have no matching SST value and are excluded from the final joined dataset). The subtitle on each panel gives the count and percentage of assemblage samples falling outside SST coverage for that core.

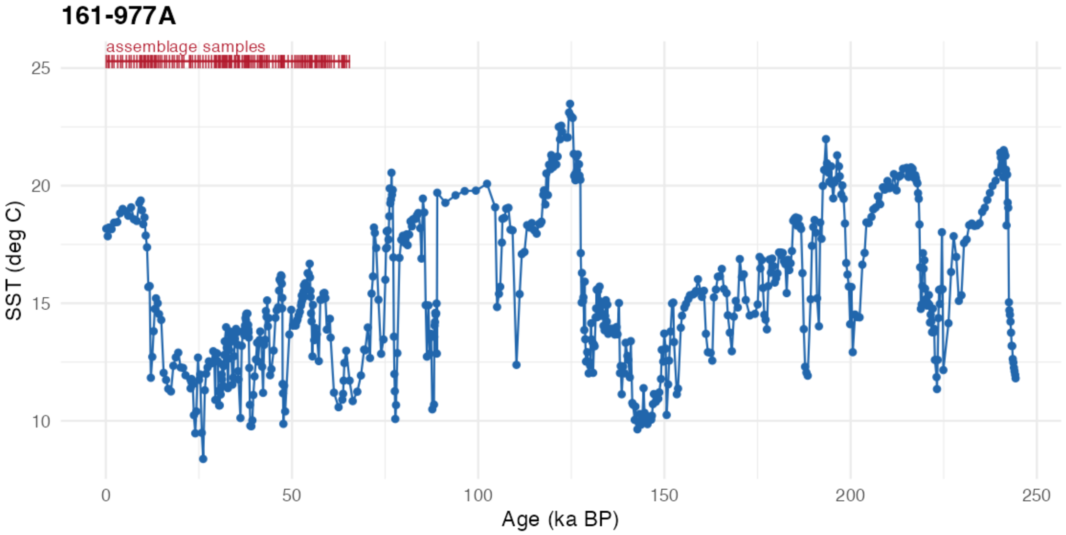


**FIGURE S9** As Fig. S8, for core 161-977A.

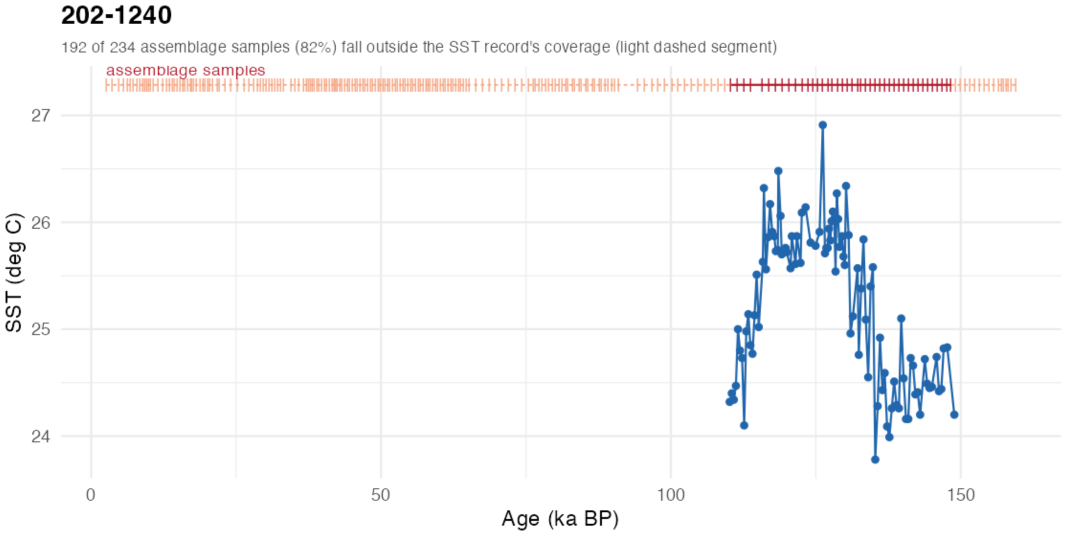


**FIGURE S10** As Fig. S8, for core 202-1240.

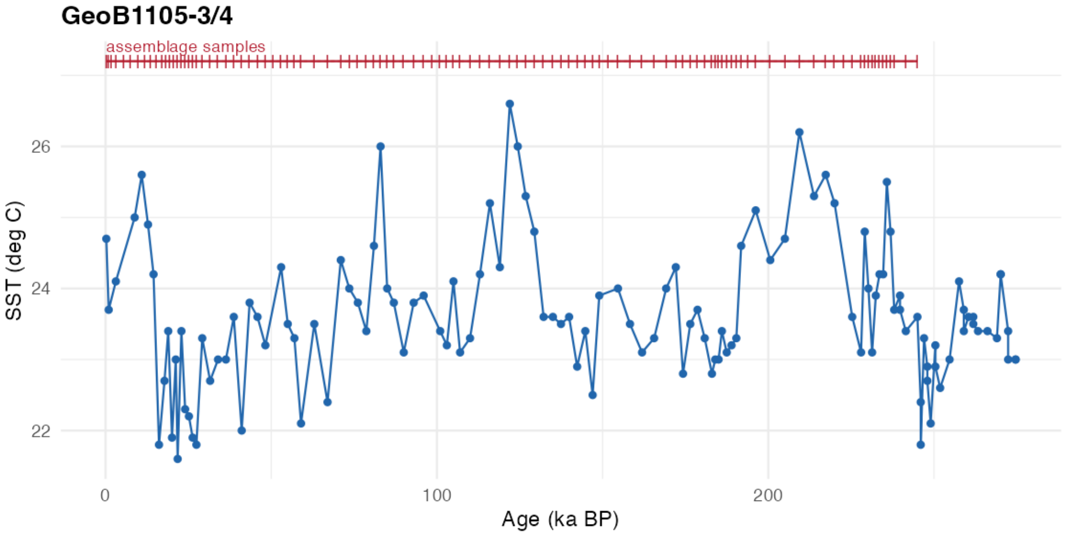


**FIGURE S11** As Fig. S8, for core GeoB1105-3/4.

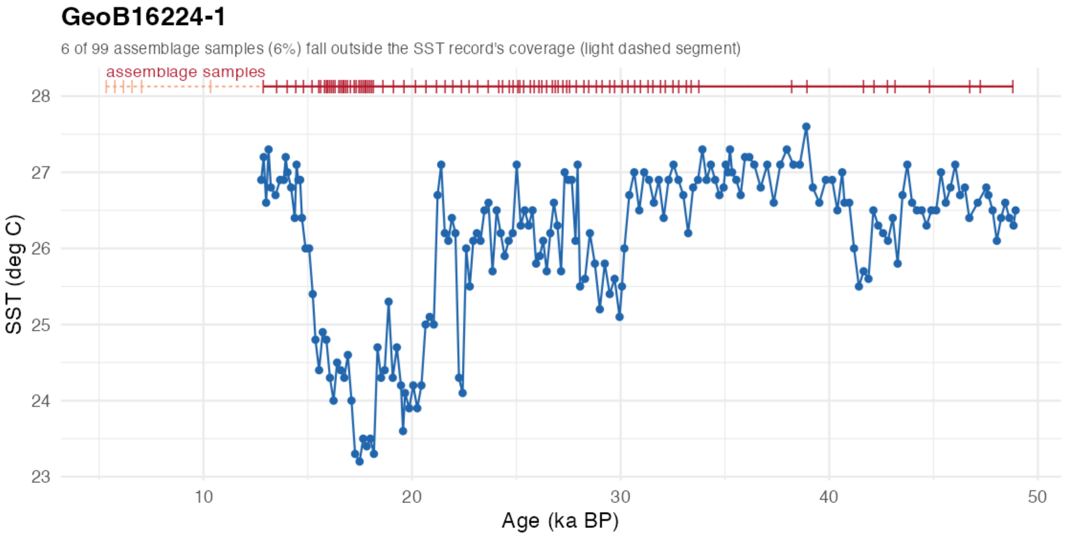


**FIGURE S12** As Fig. S8, for core GeoB16224-1.

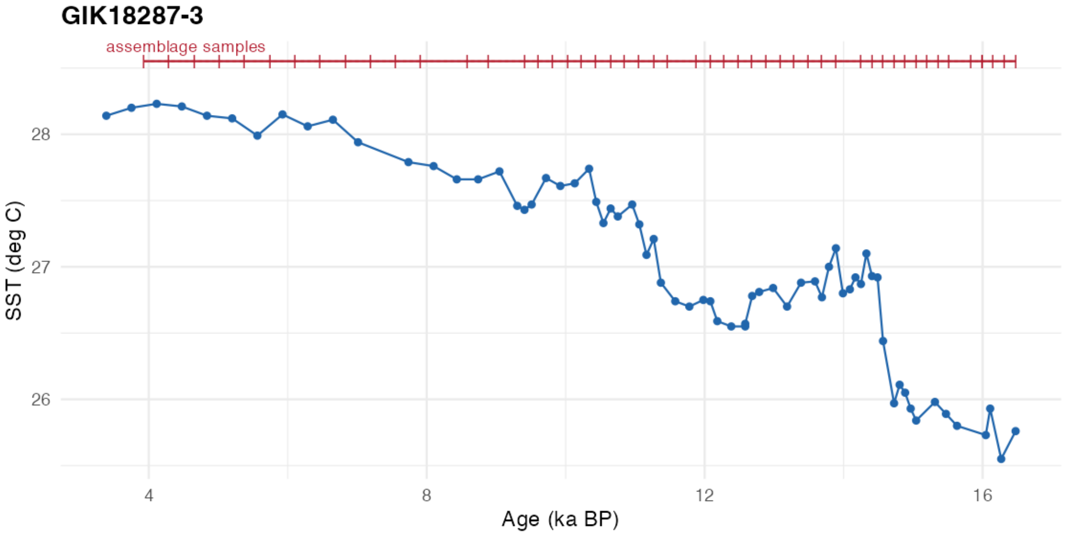


**FIGURE S13** As Fig. S8, for core GIK18287-3.

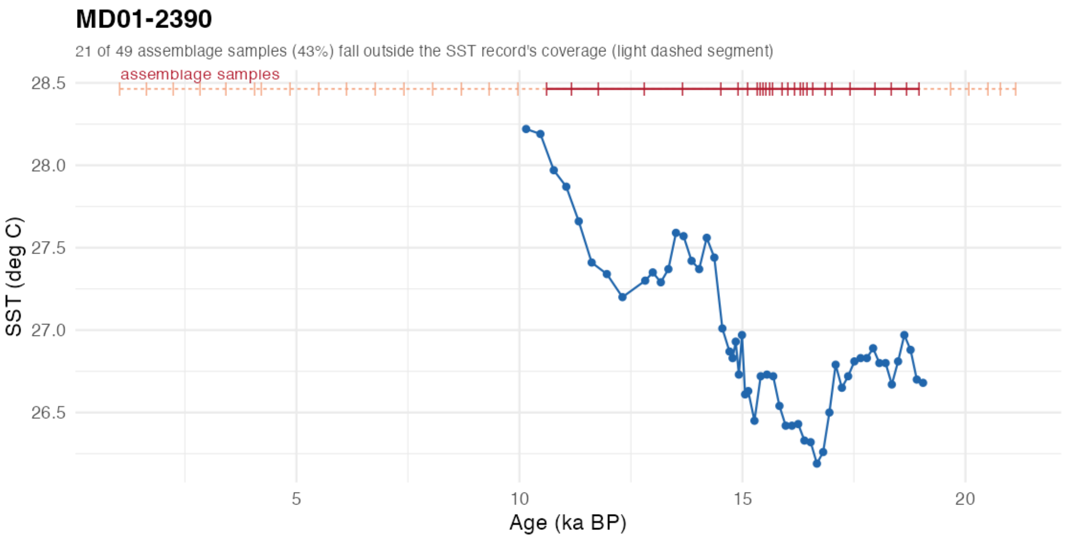


**FIGURE S14** As Fig. S8, for core MD01-2390.

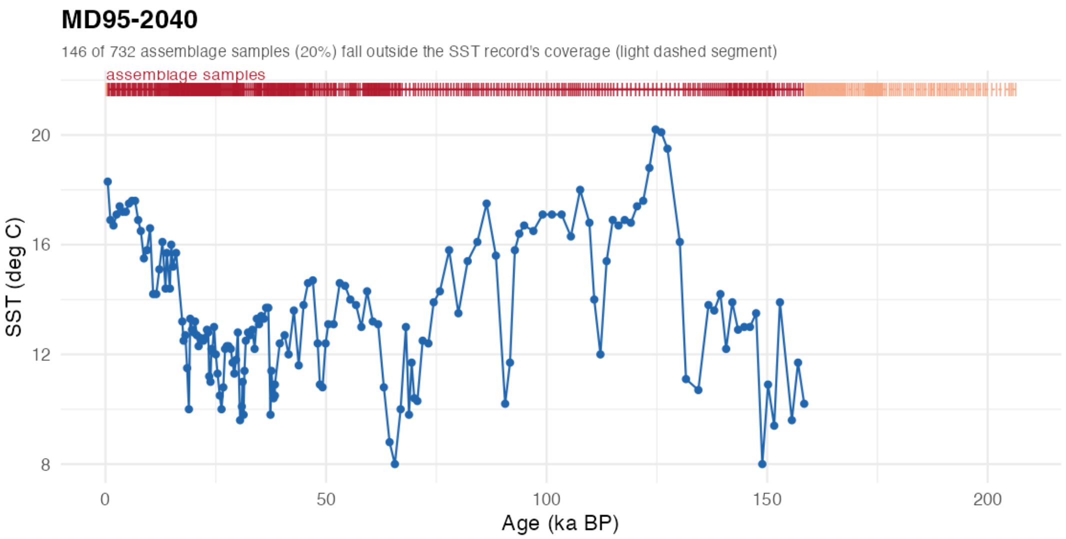


**FIGURE S15** As Fig. S8, for core MD95-2040.

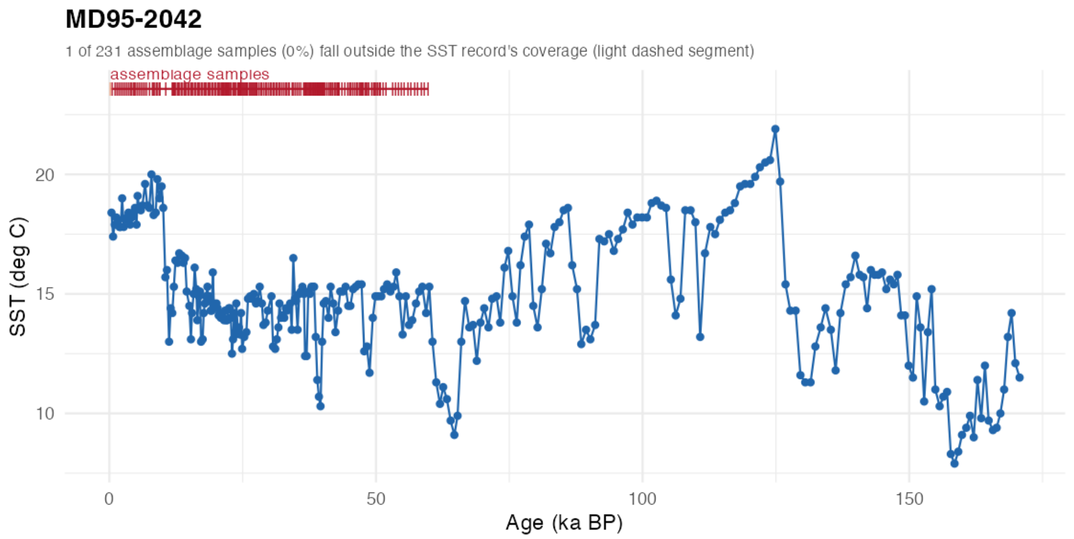


**FIGURE S16** As Fig. S8, for core MD95-2042.

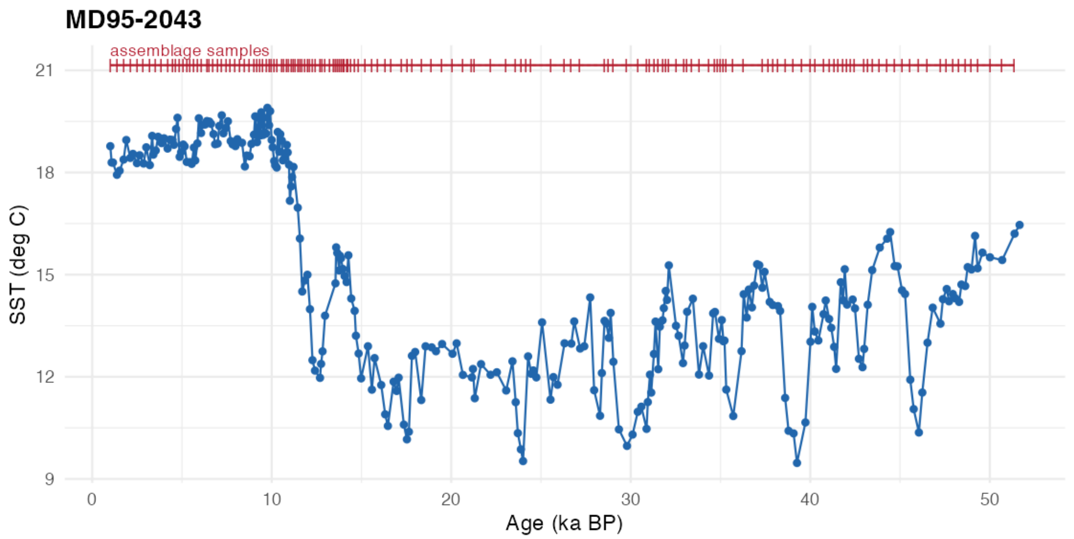


**FIGURE S17** As Fig. S8, for core MD95-2043.

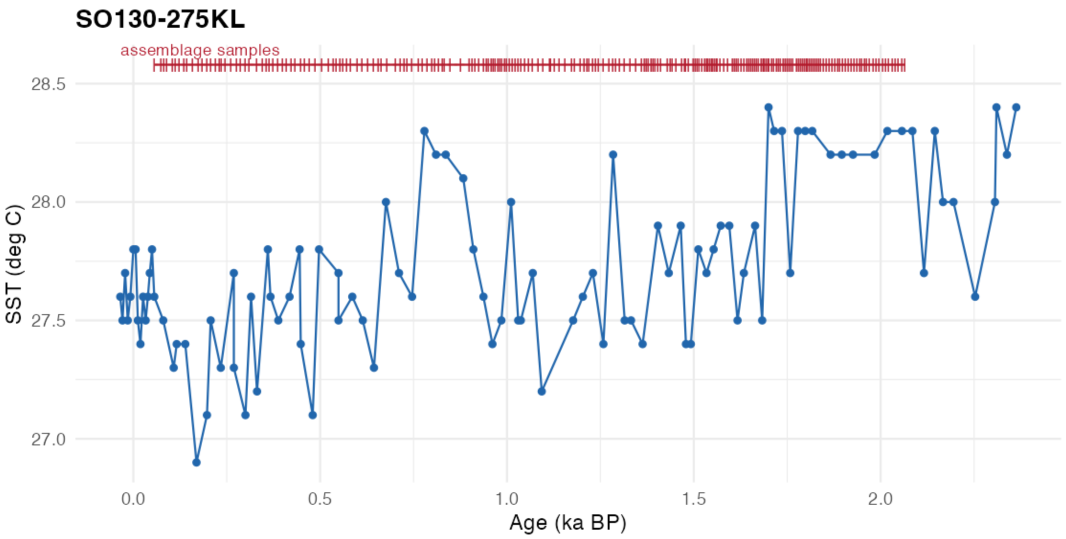


**FIGURE S18** As Fig. S8, for core SO130-275KL.

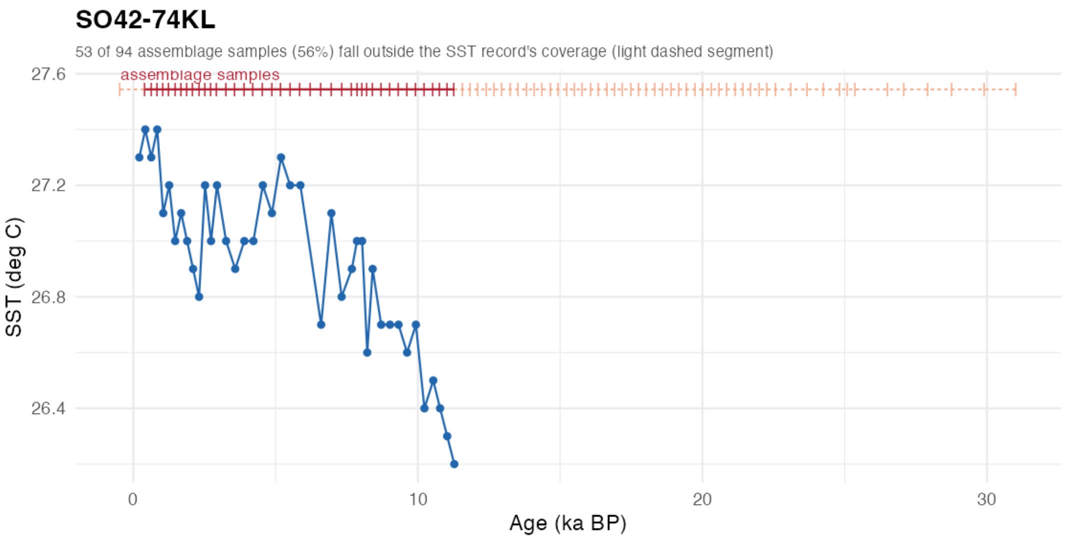


**FIGURE S19** As Fig. S8, for core SO42-74KL.

**TABLE S3** Source data used in this study. All raw data – sea-surface temperature (SST) reconstructions, planktonic foraminifera assemblage/census counts, age models – originate from previously published, DOI-archived sources; see main text for how these were standardized and combined.

| Core | Data type | Citation & DOI | License |
|---|---|---|---|
| 161-975 | SST (UK'37) | Kandiano, E.S.; Bauch, H.A.; Fahl, K. (2014): Alkenones and UK'37 derived SSTs of late MIS 6 to early MIS 5d section of ODP Site 161-975. PANGAEA. https://doi.org/10.1594/PANGAEA.836211 | CC-BY-3.0 |
| 161-975 | Assemblage | Kandiano, E.S.; Bauch, H.A.; Fahl, K. (2014): Planktonic foraminifera and SSTs of late MIS 6 to early MIS 5d section of ODP Site 161-975. PANGAEA. https://doi.org/10.1594/PANGAEA.836212 | CC-BY-3.0 |
| 161-977A | SST (UK'37) | Martrat, B.; Grimalt, J.O.; Shackleton, N.J.; de Abreu, L.; Hutterli, M.A.; Stocker, T.F. (2007): (Table S2) Sea surface temperature estimation for ODP Hole 161-977A. PANGAEA. https://doi.org/10.1594/PANGAEA.771890 | CC-BY-3.0 |
| 161-977A | Assemblage | Pérez-Folgado, M.; Sierro, F.J.; Flores, J-A.; Cacho, I.; Grimalt, J.O.; Zahn, R.; Shackleton, N.J. (2003): (Appendix 1) Distribution of planktonic foraminifera in sediment of the last 70 kyr of ODP Core 161-977A. PANGAEA. https://doi.org/10.1594/PANGAEA.678285 | CC-BY-3.0 |
| 202-1240 | SST (UK'37) | Quirós-Collazos, L.; Calvo, E.; Schouten, S.; Rodrigo-Gámiz, M.; Pena, L.D.; Cacho, I.; Pelejero, C. (2020): Molecular biomarkers and UK'37-derived sea surface temperature of ODP Site 202-1240. PANGAEA. https://doi.org/10.1594/PANGAEA.916208 | CC-BY-4.0 |
| 202-1240 | Assemblage | Yu, P-S.; Kienast, M.; Chen, M-T.; Cacho, I.; Flores, J-A.; Mohtadi, M.; Mix, A.C. (2012): Distribution of planktic foraminifera in ODP Site 202-1240 samples. PANGAEA. https://doi.org/10.1594/PANGAEA.807101 | CC-BY-3.0 |
| GIK18287-3 | SST (UK'37) | Steinke, S.; Kienast, M.; Pflaumann, U.; Weinelt, M.; Stattegger, K. (2001): UK'37 and sea surface temperatures of sediment core GIK18287-3. PANGAEA. https://doi.org/10.1594/PANGAEA.80225 | CC-BY-3.0 |

*(continued on next page)*

*(continued from previous page)*

| Core | Data type | Citation & DOI | License |
|---|---|---|---|
| GIK18287-3 | Assemblage | Steinke, S.; Kienast, M.; Pflaumann, U.; Weinelt, M.; Stattegger, K. (2001): Distribution of planktic foraminifera of sediment core GIK18287-3. PANGAEA. https://doi.org/10.1594/PANGAEA.80123 | CC-BY-3.0 |
| GeoB1105-3/4 | SST (Mg/Ca) | Nürnberg, D.; Müller, A.; Schneider, R.R. (2000): Paleo-sea surface temperatures and Mg/Ca ratios of sediment cores GeoB1105-3/4. PANGAEA. https://doi.org/10.1594/PANGAEA.710804 | CC-BY-3.0 |
| GeoB1105-3/4 | Assemblage | Wefer, G.; Berger, W.H.; Bickert, T.; et al. (1996): Distribution of planktonic foraminifera of site GeoB1105. PANGAEA. https://doi.org/10.1594/PANGAEA.54710 | CC-BY-3.0 |
| GeoB16224-1 | SST (UK'37) | Crivellari, S.; Chiessi, C.M.; Kuhnert, H.; et al. (2019): UK'37-based temperature record of sediment core GeoB16224-1. PANGAEA. https://doi.org/10.1594/PANGAEA.902159 | CC-BY-4.0 |
| GeoB16224-1 | SST (Mg/Ca) | Crivellari, S.; Chiessi, C.M.; Kuhnert, H.; et al. (2019): Mg/Ca-based temperature record of sediment core GeoB16224-1. PANGAEA. https://doi.org/10.1594/PANGAEA.902154 | CC-BY-4.0 |
| GeoB16224-1 | SST (TEX) | Crivellari, S.; Chiessi, C.M.; Kuhnert, H.; et al. (2019): TEX86H and TEX86L temperature records of sediment core GeoB16224-1. PANGAEA. https://doi.org/10.1594/PANGAEA.902160 | CC-BY-4.0 |
| GeoB16224-1 | Assemblage | No public/DOI, provided by Rodrigo Costa Portilho-Ramos; related publication: Crivellari et al. 2019, *EPSL*, https://doi.org/10.1016/j.epsl.2019.05.006. | N/A |
| MD01-2390 | SST (UK'37) | Steinke, S. (2008): Alkenon-based sea surface temperatures (annual and winter SST) of sediment core MD01-2390. PANGAEA. https://doi.org/10.1594/PANGAEA.690520 | CC-BY-3.0 |
| MD01-2390 | Assemblage | Steinke, S.; Yu, P-S.; Kucera, M.; Chen, M-T. (2008): (Appendix Table 3) Relative abundances of planktonic foraminiferal species in sediment core MD01-2390. PANGAEA. https://doi.org/10.1594/PANGAEA.775185 | CC-BY-3.0 |
| MD95-2040 | SST (UK'37) | Pailler, D.; Bard, E. (2002): (Table 2) Geochemical analytical data for sediment core MD95-2040. PANGAEA. https://doi.org/10.1594/PANGAEA.96865 | CC-BY-3.0 |

*(continued on next page)*

*(continued from previous page)*

| Core | Data type | Citation & DOI | License |
|---|---|---|---|
| MD95-2040 | Age model | Voelker, A.H.L.; de Abreu, L. (2011): Age model of sediment core MD95-2040. PANGAEA. https://doi.org/10.1594/PANGAEA.737133 | CC-BY-3.0 |
| MD95-2040 | Assemblage | de Abreu, L.; Shackleton, N.J.; Schönfeld, J.; Hall, M.A.; Chapman, M.R. (2003): Planktic foraminifera counts of sediment core MD95-2040. PANGAEA. https://doi.org/10.1594/PANGAEA.66714 | CC-BY-3.0 |
| MD95-2042 | SST (UK'37) | Pailler, D.; Bard, E. (2002): (Table 1) Geochemical analytical data for sediment core MD95-2042. PANGAEA. https://doi.org/10.1594/PANGAEA.96864 | CC-BY-3.0 |
| MD95-2042 | Assemblage | Rossignol, L. (2022): Planktonic foraminifera counts of sediment core MD95-2042 off the Iberian margin for the past 60 ka. PANGAEA. https://doi.org/10.1594/PANGAEA.946601 | CC-BY-4.0 |
| MD95-2043 | SST (UK'37) | Martrat, B.; Jimenez-Amat, P.; Zahn, R.; Grimalt, J.O. (2014): Sea surface temperatures, alkenones and sedimentation rates at the Alboran basin from sediment core MD95-2043. PANGAEA. https://doi.org/10.1594/PANGAEA.787985 | CC-BY-3.0 |
| MD95-2043 | Assemblage | Pérez-Folgado, M.; Sierro, F.J.; Flores, J-A.; Cacho, I.; Grimalt, J.O.; Zahn, R.; Shackleton, N.J. (2003): (Appendix 2) Distribution of planktonic foraminifera in the last 70 kyr of sediment core MD95-2043. PANGAEA. https://doi.org/10.1594/PANGAEA.678286 | CC-BY-3.0 |
| SO42-74KL | SST (UK'37) | Kim, J-H.; Rimbu, N.; Lorenz, S.J.; et al. (2004): Age and alkenone-derived Holocene sea-surface temperature records of sediment core SO42-74KL. PANGAEA. https://doi.org/10.1594/PANGAEA.438818 | CC-BY-3.0 |
| SO42-74KL | Assemblage | Schulz, H. (1995): Planktic foraminifera assemblage in sediment core SO42-74KL. PANGAEA. https://doi.org/10.1594/PANGAEA.134146 | CC-BY-3.0 |
| SO130-275KL | SST (UK'37) | Böll, A.; Lückge, A.; Munz, P.; Forke, S.; Schulz, H.; Ramaswamy, V.; Rixen, T.; Gaye, B.; Emeis, K-C. (2014): Sea surface temperature reconstruction for sediment cores SO90-39KG and SO130_275KL. PANGAEA. https://doi.org/10.1594/PANGAEA.839062 | CC-BY-3.0 |

*(continued on next page)*

*(continued from previous page)*

| Core | Data type | Citation & DOI | License |
|---|---|---|---|
| SO130-275KL | Assemblage | Munz, P.; Siccha, M.; Lückge, A.; Böll, A.; Kucera, M.; Schulz, H. (2015): Distribution of planktic foraminifera of sediment core SO130-275KL. PANGAEA. https://doi.org/10.1594/PANGAEA.853967 | CC-BY-3.0 |
